# Climate change threatens the productivity stability of Italian forests

**DOI:** 10.64898/2026.09.30.755743

**Authors:** Mauro Morichetti, Daniela Dalmonech, Elisa Grieco, Elia Vangi, Alessio Collalti

## Abstract

Mediterranean forests are emerging as one of the clearest hotspots of climate-change impacts, where rapid warming and increasingly severe droughts and heatwaves are pushing forest ecosystems toward their physiological and functional limits. These pressures threaten productivity, carbon dynamics, ecosystem stability, and the services forests provide. In this context, a process-based forest ecosystem model was used to assess climate-driven changes in carbon dynamics and productivity stability across Italian forests within the Inter-Sectoral Impact Model Intercomparison Project (ISIMIP) framework. 5135 plots were simulated, corresponding to ∼80% of the field plots included in the second Italian National Forest Inventory (INFI). Model simulations were first evaluated against independent observations of gross primary production (GPP) from satellite-based products and forest carbon variables from the third INFI. Forest dynamics from 2015 to 2100 were simulated under three CMIP6-based emission scenarios (SSP1-2.6, SSP3-7.0, and SSP5-8.5) using two climate forcings (GFDL-ESM4 and UKESM1-0-LL). Future responses were assessed for GPP, aboveground biomass, growing stock volume, and carbon stocks, while variability in GPP was quantified using the coefficient of variation as an indicator of forest stability. Results show that the model reproduced the main spatial and temporal patterns of forest productivity and carbon stocks. Future projections revealed strong spatial heterogeneity and increasing divergence among scenarios after mid-century. GPP showed the earliest and strongest response to climate change, with mountain forests generally maintaining or increasing productivity and exhibiting greater stability, whereas coastal forests showed pronounced productivity declines and increasing variability, particularly under higher-emission scenarios and more extreme climate forcing. In contrast, carbon stocks generally continued to increase throughout the century, although carbon accumulation progressively slowed toward the end of the century, particularly under higher-emission scenarios.

## 1. Introduction

The Mediterranean has been identified as one of the regions most responsive to global climate change, with observed warming expected to continue at a rate exceeding the global average (FAO, 2019; Giorgi and Bi, 2005; Lionello and Scarascia, 2018; Noce et al., 2016). Warming, drying, and increasing evaporative demand are intensifying drought stress and exacerbating the existing water limitations on vegetation (Gouveia et al., 2017; Lazoglou et al., 2024; Tramblay et al., 2020). These changes are particularly concerning for forests, as temperature and water availability directly regulate key physiological processes, including photosynthesis, respiration, growth, and carbon allocation, while increasingly frequent compound hot-dry events can further increase tree mortality and disturbance risk (Grünig et al., 2026). Across Europe, excess forest mortality rises sharply once integrated spring-summer water balance crosses a threshold, and drought caused about 500,000 ha of excess mortality from 1987 to 2016 (Senf et al., 2020). Compound hot-dry events are especially important: 46% of documented European mortality events coincided with rare hot-summer plus dry-year combinations, and these events have become increasingly common since the 1980s (Gazol and Camarero, 2022). Droughts in the Mediterranean region are expected to increase in frequency and severity under future climate change scenarios, but future uncertainties remain, highlighting the need to maintain forest stability and strengthen adaptive capacity (FAO 2019; Giorgi and Bi, 2005). Understanding how forests will respond to future climate conditions is therefore essential for assessing their future role in climate mitigation and for supporting effective adaptation (Barbati et al., 2018).

However, Mediterranean forests do not exhibit a uniform response to climate change, with their sensitivity varying substantially along gradients of climate, topography, elevation, soil conditions, species traits, and stand characteristics (Carnicer et al., 2021; Dorado-Liñán et al., 2019; FAO, 2019; Puchi et al., 2024; Santini et al., 2014; Miglietta & Valentini, 2015). Warming may partially alleviate temperature limitation in colder, high-elevation environments, whereas increasing temperature and atmospheric water demand can intensify water limitation in warmer and drier areas, resulting in contrasting responses ranging from growth increases in cold mountain forests to productivity declines, turnover, and dieback in warmer and more water-limited sites (Charru et al., 2017; Colangelo et al., 2021; Dorado-Liñán et al., 2019; Fibbi et al., 2019). Italy provides a particularly suitable domain for investigating these processes, as its forests encompass pronounced gradients in climate, elevation, water availability, and vegetation composition. The Alpine and Apennine mountain systems, together with the Po Plain and extensive Mediterranean coastal areas, support forest ecosystems ranging from cold-adapted conifers to drought-adapted Mediterranean evergreen forests (Conte et al., 2019; Vacchiano et al., 2017). These environmental gradients expose Italian forests to contrasting climatic constraints, with drought emerging as an important driver of productivity declines, although its effects depend on species, provenance, and forest type (Fibbi et al., 2019; Nolè et al., 2013; Rita et al., 2020). Evidence also indicates that climate-related stress can influence Italian forests through changes in growth, mortality, and stand dynamics (Hillebrand et al., 2023; Noce et al., 2023).

Gross primary production (GPP) is the largest terrestrial carbon flux and a key indicator of forest ecosystem functioning and carbon uptake (Beer et al., 2010). Even relatively small changes in GPP have shown to substantially affect the terrestrial carbon balance and, consequently, the exchange of CO₂ between ecosystems and the atmosphere (O’Sullivan et al., 2020; Yao et al., 2018; Zheng et al., 2020). Because photosynthesis is directly regulated by temperature and water availability, GPP is highly responsive to climatic variability, and changes in GPP can therefore provide an early indication of climate-driven changes in ecosystem functioning, particularly under extreme temperatures and precipitation deficits (Xu et al., 2024; You et al., 2020; Zhu et al., 2026). However, annual GPP alone may mask changes in ecosystem functioning, as forests can maintain similar productivity levels while experiencing increasing year-to-year variability (Lin et al., 2023; Zhang et al., 2022). The variability in primary productivity therefore provides complementary information on the temporal stability of ecosystem functioning. However, assessing changes in forest functioning requires considering not only the dynamics of carbon uptake, but also how carbon is accumulated and retained within forest biomass over time. Carbon stocks provide this longer-term perspective, as they integrate the cumulative effects of carbon assimilation, allocation, growth, and disturbance. In this study, aboveground biomass (AGB), growing stock volume (GSV), and carbon stocks (CS) alongside GPP and its variability were considered to capture complementary dimensions of forest carbon dynamics, from short-term ecosystem functioning to longer-term carbon storage.

Process-based models (PBMs) provide a useful basis for investigating how climate change may alter forest carbon dynamics, including both carbon fluxes (i.e., GPP), and carbon stocks, (i.e., AGB, GSV, and CS) (Maréchaux et al., 2021; Vacchiano et al., 2012). At the national scale in Italy, Vangi et al. (2025) applied the process-based forest model 3D-CMCC-FEM (Collalti et al., 2024) to simulate forest carbon fluxes and stocks under current climate. Model predictions were evaluated against remote sensing-derived datasets and observations from the third Italian National Forest Inventory (INFI; INFC, 2021), showing a good agreement with the observed spatial and temporal patterns of forest carbon dynamics. Building on these findings, we applied and evaluated a biogeochemical, biophysical, process-based forest ecosystem model (namely 3D-CMCC-FEM) under modelled historical climate conditions and subsequently used the model to investigate forest carbon dynamics under future climate scenarios. The simulations were performed on a number of forest stands distributed across Italy and initialized using structural and compositional information from the second INFI (INFC, 2007) following closely the framework simulation from Vangi et al. (2025). Future simulations were conducted within the Inter-Sectoral Impact Model Intercomparison Project (ISIMIP; Reyer et al., 2019, 2020) context, which provides harmonized, bias-adjusted, and statistically downscaled CMIP6 climate forcing for consistent impact assessments. We considered three socioeconomic pathways representing progressively increasing radiative forcing, SSP1-2.6, SSP3-7.0, and SSP5-8.5, and used two Earth System Models (ESMs), GFDL-ESM4 and UKESM1-0-LL, to account for differences in projected climate conditions.

Specifically, we address the following questions: (i) whether 3D-CMCC-FEM, driven by three historical climate forcings (i.e., ERA5-Land, GFDL-ESM4, and UKESM1-0-LL), cans adequately represent forest carbon dynamics under modelled current climate conditions by evaluating simulated GPP against independent observations from remote-sensing products, and simulated AGB, GSV, and CS against observations from the INFI 2015; (ii) how, to what extent, and by how much forest carbon fluxes and stocks are projected to change under alternative future climate conditions within the ISIMIP framework; and (iii) whether the variability of GPP changes under future climate conditions and whether these changes differ across the ecoregions represented by Italian forests, thereby providing insights into the temporal stability of forest productivity.

## 2. Materials and methods

### 2.1. Italian case study

Italy covers approximately 301,000 km², extending along a northwest-southeast axis across the Mediterranean Sea and spanning ∼35° to 47° N (Fig. 1). Its broad latitudinal range, elevations from sea level to over 4,000 m, and approximately 7,500 km of coastline generate marked climatic and environmental heterogeneity. The landscape is dominated by the Alps and Apennines, with major lowlands concentrated in the Po Plain and along the coasts. This geographical diversity supports forest ecosystems ranging from Alpine coniferous forests to Mediterranean evergreen woodlands (Alberico et al., 2023; Vangi et al., 2025b). To investigate the regional variability of climate change impacts, forest stands were classified according to the ecoregional framework proposed by Blasi et al. (2018). The Italian ecoregional classification accounts for differences in macroclimate, physiography, and potential natural vegetation, thereby capturing major bioclimatic, orographic, and biogeographical gradients across the country. In this study, the province level was selected as the spatial unit for the analysis, since it captures major environmental and vegetation gradients while providing a suitable spatial scale for national forest assessments. The analysis included five of Italy’s seven ecoregional provinces: Alpine, Po Plain, Apennine, Tyrrhenian, and Adriatic (Fig. 1). The Alpine and Apennine provinces encompass the main temperate and mountainous sectors of Italy, whereas the Po Plain, Tyrrhenian and Adriatic provinces capture distinct coastal and Mediterranean environments (Blasi et al., 2014; Gatti et al., 2024).

**Figure 1:**
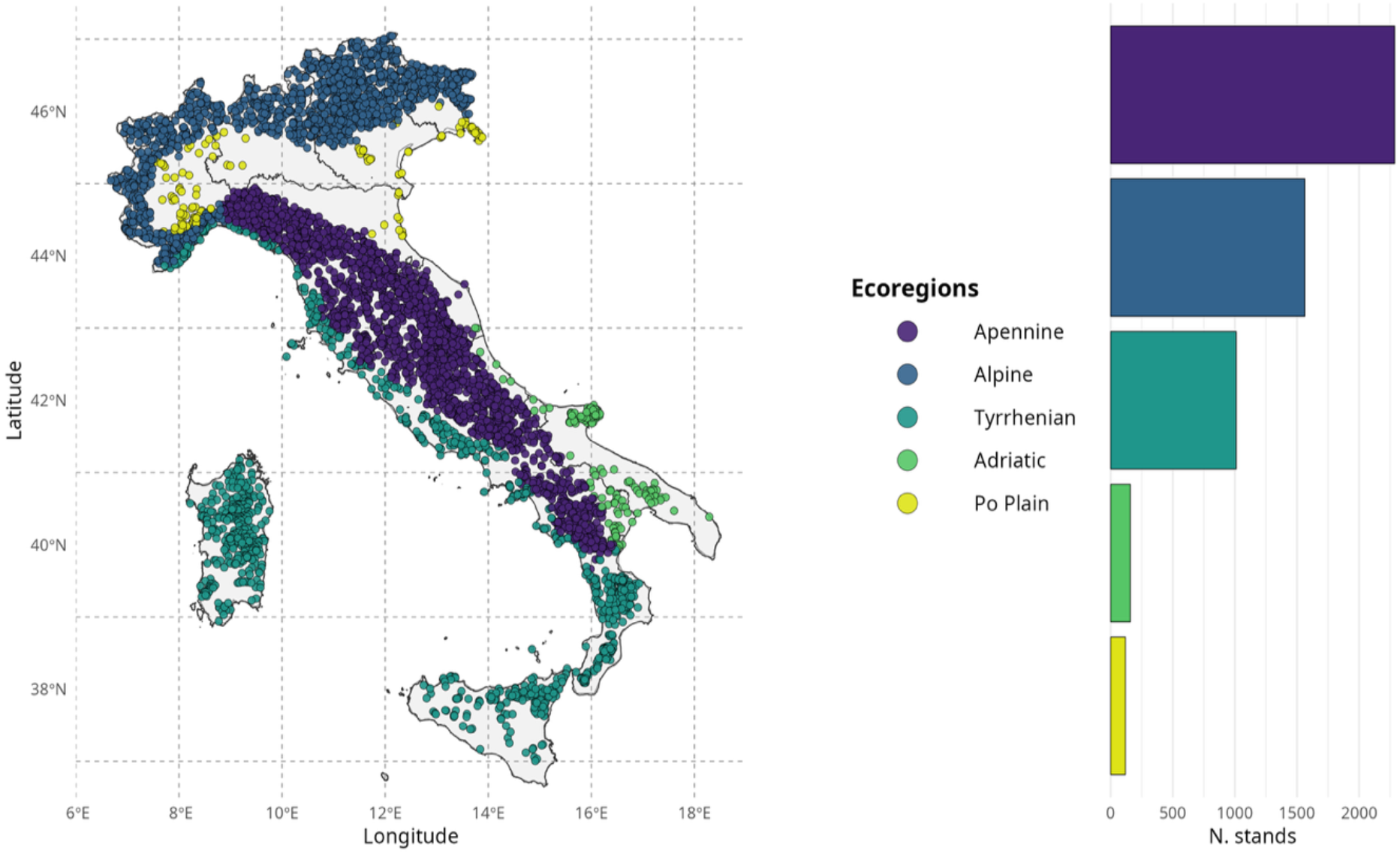
Spatial distribution of forest stands across Italy classified by ecoregions. Each point represents a modelled stand and is colored according to its ecoregion: Alpine (blue), Apennine (purple), Tyrrhenian (teal), Adriatic (green), and Po Plain (yellow). The bar chart on the right shows the number of stands per ecoregion.

According to the third INFI, forests and other wooded lands cover approximately 11 million hectares, corresponding to about 36% of the national territory (INFC, 2021). As shown in Figure 2, ‘*Mixed*’ forests represented the largest proportion of the simulated stands and were distributed across all Italian ecoregions, providing a broad representation of the structural and ecological diversity of Italian forests is reflected in their dominant forest types. Alpine forests were primarily characterized by *Picea abies* (L.) H. Karst. and *Larix decidua* Mill., whereas the Apennine ecoregion was dominated by *Fagus sylvatica* L. and *Quercus cerris* L. Forest types typical of the coastal ecoregions, including *Quercus ilex* L., *Pinus halepensis* Mill., and *Pinus nigra* subsp. *laricio* (Poir.) Maire were mainly associated with the Tyrrhenian and Adriatic ecoregions, while *Quercus robur* L. was representative of the Po Plain (Vangi et al., 2025b).

**Figure 2:**
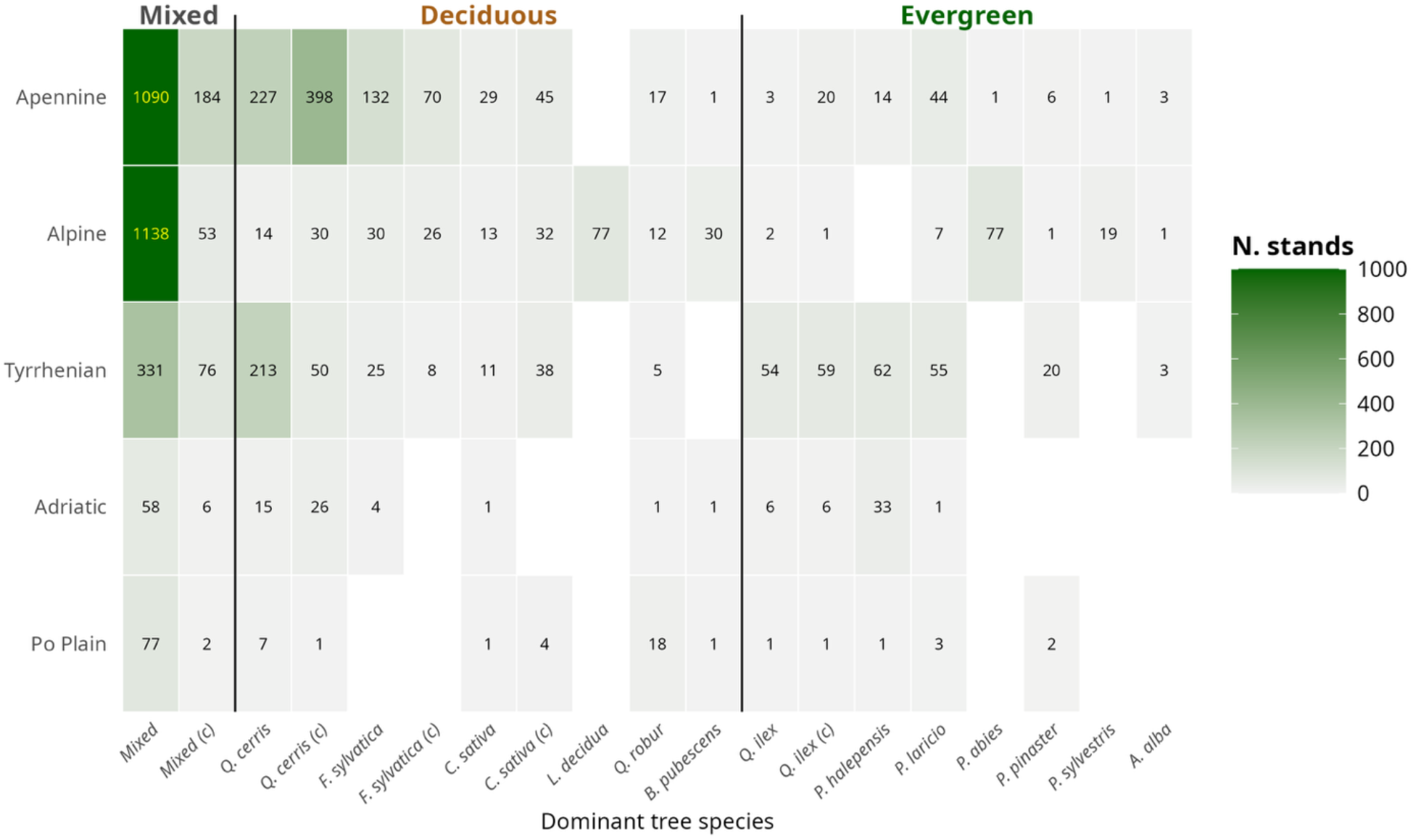
Distribution of forest stands by dominant tree species and ecoregion. Rows represent ecoregions (i.e., Apennine, Alpine, Tyrrhenian, Adriatic, and Po Plain), while columns report dominant species. Species are grouped into mixed (left), deciduous (center) and evergreen (right) functional types. Values and color intensity indicate the number of stands (N. stands) associated with each species-ecoregion combination. The suffix (c) indicates coppice stands.

### 2.2. The 3D-CMCC-FEM model

#### 2.2.1. Model description

Forest growth and carbon dynamics were simulated using the “Three Dimensional – Coupled Model Carbon Cycle – Forest Ecosystem Module” (namely, 3D-CMCC-FEM, version 5.7), a process-based forest ecosystem model that simulates the dynamics of carbon, nitrogen, and water cycles under changing environmental conditions, forest management, and climate scenarios (Collalti et al., 2016, 2018, 2022, 2024; Dalmonech et al., 2022; Vangi et al., 2025a). The model represents main eco-physiological processes at the stand scale (i.e., 1 ha), while individual processes are simulated at different temporal resolutions, ranging from daily to monthly or annual time steps. More recently, 3D-CMCC-FEM has been successfully applied first at the regional scale and subsequently at the national scale (Dalmonech et al., 2026, 2024; Vangi et al., 2025b). The 3D-CMCC-FEM has been extensively evaluated across varied forest ecosystems and climatic conditions and applied to investigate forest productivity, carbon sequestration, forest management, and ecosystem responses to climate change (Collalti et al., 2018, 2016; Mahnken et al., 2022; Marconi et al., 2017; Morichetti et al., 2024; Puchi et al., 2026; Saponaro et al., 2026, 2025; Testolin et al., 2023; Vangi et al., 2024a, 2024b). Further details on the model description are provided in the Supplementary Material (Section 1.1.1, Model description).

The 3D-CMCC-FEM requires input data describing the forest stand conditions at the, including species composition, diameter at breast height (AvDBH), tree height (Height), stand age (Age), tree density (N), and soil properties (CLAY_PERC, SILT_PERC, SAND_PERC and SOIL_DEPTH). Daily meteorological forcing includes average (Ta_f), maximum (Tmax) and minimum (Tmin) air temperature, relative humidity (RH_f), downward shortwave solar radiation (Rg_f), and precipitation (Precip), together with atmospheric CO₂ concentration (Collalti et al., 2022). The model also needs species-specific parameters describing eco-physiological and biophysical attributes, which are derived mainly from the literature and previous model validation studies, allowing the model to be applied across a range of forest ecosystems while minimizing the need for site-specific calibration (Mahnken et al., 2022).

As outputs, the model provides a wide range of variables at daily, monthly and annual temporal resolution, describing forest carbon, nitrogen, water and energy dynamics, as well as stand structure and productivity. In this study, five key variables were considered: GPP, AGB, belowground biomass (BGB), GSV, and CS. GPP, AGB and BGB were directly obtained from the model outputs. AGB (tC ha⁻¹) is calculated within the model as the sum of carbon contained in leaves, stems, branches and fruits, whereas BGB (tC ha⁻¹) is calculated as the sum of fine- and coarse-root carbon pools (Collalti et al., 2022). For the present analysis, AGB was converted from carbon to dry matter (DM) using a carbon fraction of 0.5, resulting in a factor of two:

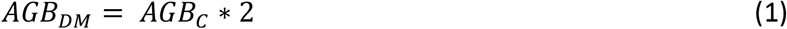

from this point onward, AGB refers to aboveground dry biomass (AGB_DM_) and is expressed in Mg ha⁻¹. GSV (m⁻^3^ ha⁻^1^) was subsequently derived from dry-matter AGB following Federici et al. (2008) using the species-specific biomass expansion factor (BEF) and wood basic density (WBD):

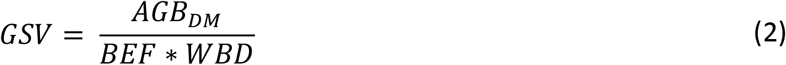

CS (MgC ha⁻^1^) was calculated directly from the carbon-based above- and belowground biomass pools, representing thus the total tree carbon stocks:

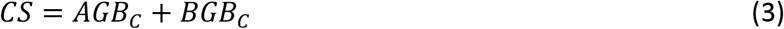

All variables were extracted at the stand level and subsequently aggregated according to the spatial and temporal scales described below.

#### 2.2.2. NFI stand initialization

Stand initialization was based on the Italian National Forest Inventory, using tree-level information collected during the second INFI field campaign at their geo-referenced locations to characterize the initial stand structure and species composition (INFC, 2007). Following the stand selection and initialization procedure described already by (Vangi et al., 2025b), 5135 of the 6174 available plots (∼83%) were selected for simulation, encompassing 14 of the most common Italian tree species. The selected plots included both pure and mixed-species stands (Figure 2). Stands in which more than one tree species was present were considered *‘Mixed’* stands, regardless of the relative basal-area contribution of the individual species. Simulations were initialized in 2005 and performed at a representative scale of 1 ha, with all model outputs referring to the corresponding simulation grid cell size. The simulations covered the period 2005-2100 and were conducted under a no-management scenario, with no thinning or harvesting operations applied throughout the simulation period (Vangi et al., 2025b) to highlight the purely climate impact on forest ecosystem dynamics.

#### 2.2.3. Driven climate forcing

Meteorological forcing was obtained following the ISIMIP framework (Reyer et al., 2019, 2020). The ISIMIP3b version, applied in this study, uses climate simulations from different ESMs, which are bias-adjusted and statistically downscaled using the ISIMIP3BASD methodology (Lange, 2019; Lange et al., 2020). This procedure adjusts the statistical characteristics of the simulated climate while preserving the temporal trends of the original climate simulations. Bias adjustment and statistical downscaling are performed against the WFDE5 (WATCH Forcing Data methodology applied to ERA5) and ERA5 (Fifth-generation ECMWF Reanalysis) observational reference datasets, which together form the W5E5 (WFDE5 over land and ERA5 over the ocean) dataset (Cucchi et al., 2020). The resulting ISIMIP3b forcing is provided at a spatial resolution of 0.5° (∼50 km) and a daily temporal resolution, covering historical (1850-2014) and future periods (2015-2100) globally.

In the present study, two ISIMIP3b ESMs, GFDL-ESM4 and UKESM1-0-LL, were selected to provide alternative projections of future climate conditions (Dunne et al., 2020; Sellar et al., 2020). Both models are part of the Coupled Model Intercomparison Project Phase 6 (CMIP6) (Eyring et al., 2016). Future CMIP6 projections are organized within the Scenario Model Intercomparison Project (ScenarioMIP), which provides climate simulations under alternative socioeconomic and greenhouse-gas emission pathways. The three scenarios considered here were SSP1-2.6, SSP3-7.0, and SSP5-8.5 (hereafter SSP126, SSP370 and SSP585), representing low (2.6 W m⁻²), intermediate-to-high (7.0 W m⁻²), and high future radiative-forcing (8.5 W m⁻²) pathways, respectively (Meinshausen et al., 2020; O’Neill et al., 2016). Future projections covered 2015-2100 for both ESMs and all three SSPs, resulting in six ESM-scenario combinations that were applied consistently across all simulated forest stands. For the historical period, a common simulation period of 2005-2014 was considered across the climate forcings considered. During this period, 3D-CMCC-FEM was driven by three alternative climate forcings: ERA5-Land, GFDL-ESM4, and UKESM1-0-LL. ERA5-Land was used as a monitoring-based reference, whereas GFDL-ESM4 and UKESM1-0-LL provided the corresponding ESM-based historical climate forcings. ERA5-Land is a land-surface reanalysis produced by ECMWF within the Copernicus Climate Change Service. It is generated by replaying the land component of ERA5 using downscaled meteorological forcing and has a spatial resolution of 0.1° (∼9 km) and an hourly temporal resolution (Dalmonech et al., 2026; Muñoz-Sabater et al., 2021)(Table S1).

#### 2.2.4. Soil data

Within 3D-CMCC-FEM, soil depth and texture contribute to the representation of soil water availability. The model uses a single-layer bucket representation of the soil water balance, in which available soil water is updated daily according to water inputs and losses. Soil depth determines the effective soil volume available for water storage and uptake, while soil texture contributes to the estimation of soil water-holding characteristics. Consequently, spatial differences in soil properties influence plant water availability and can affect photosynthesis through stomatal regulation, and forest carbon allocation dynamics (Collalti et al., 2024, 2016; Dalmonech et al., 2024; Saponaro et al., 2025). In the present study, soil properties were obtained from the national maps developed by the Soil Cartography Laboratory of the Council for Agricultural Research and Economics (CREA) (Corona et al., 2023; Costantini and Dazzi, 2013). The dataset consists of four layers at 250 m spatial resolution, describing the weighted average content of silt, sand and clay (%) in the upper 1 m of soil across Italy. Soil properties were spatially extracted at the location of each simulated stand, following the national-scale approach described by (Vangi et al., 2025b).

### 2.3. Experimental design

#### 2.3.1. Model evaluation

The model evaluation was conducted consistently across the three historical climate forcings (i.e., ERA5-Land, GFDL-ESM4, and UKESM1-0-LL) and the five Italian ecoregions (i.e., Adriatic, Alpine, Apennine, Po Plain, and Tyrrhenian). Two complementary satellite-based GPP products were used: GOSIF (Li and Xiao, 2019a, 2019b) and MODIS (Running and Zhao, 2021). For both products, annual GPP estimates were matched to the corresponding simulated stands and compared with 3D-CMCC-FEM outputs for 2005-2014 period. Model performance was also evaluated against the INFC 2015 (INFC, 2021). The modelled forest state corresponding to 2015 was compared with inventory observations for three variables: GSV, AGB, and CS (Vangi et al., 2025b). Model-observation agreement was quantified using five complementary statistical metrics: bias, mean absolute error (MAE), root mean square error (RMSE), Pearson’s correlation coefficient (r), and the coefficient of determination (R²). Further details on the experimental design for model evaluation are provided in the Supplementary Material (Section 1.2.1., Model evaluation).

#### 2.3.2. Analysis of changes under different climate scenarios

The analysis of changes under different climate scenarios followed the same spatial methodology adopted for model evaluation. Model outputs were spatially aggregated using the five Italians ecoregions as the spatial units of the analysis (i.e., Adriatic, Alpine, Apennine, Po Plain, and Tyrrhenian). In this part of the study, the future simulations (2015-2100) were analyzed for the three climate scenarios (i.e., SSP126, SSP370, and SSP585) driven by the two ESMs (i.e., GFDL-ESM4 and UKESM1-0-LL). Annual model outputs were computed over 10-year periods to reduce the influence of individual years and facilitate comparison among periods and scenarios. Each period was labelled using the central year of the corresponding interval (e.g., 2015-2024 was labelled 2020, 2025-2034 as 2030 etc.). This approach was applied to GPP, AGB, GSV, and CS to describe their temporal evolution throughout the simulation period.

The coefficient of variation (CV) of GPP, calculated as the standard deviation of annual GPP relative to its mean, was used as an indicator of the temporal stability of forest productivity, by quantifying its relative interannual variability. CV normalizes year-to-year fluctuations by mean productivity, allowing comparisons across ecoregions, forest types, and forcing climate more interpretable. Lower CV values indicate lower variability and, therefore, greater stability of ecosystem functioning, although increasing CV indicates greater variability and lower stability (Amantai et al., 2023; Marzolf et al., 2024; Musavi et al., 2017; Smith et al., 2022; Yan et al., 2023; Zhang et al., 2022). In this study, CV was calculated for each stand and decade as the ratio between the standard deviation and mean annual GPP, expressed as a percentage:

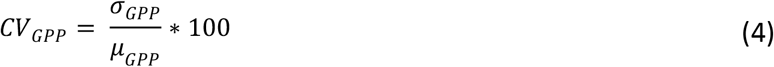

where *σ_GPP_* is the standard deviation and *μ_GPP_* is the mean annual GPP within the corresponding period. The resulting CV (%) was used to compare variability in productivity across ecoregions and ESM-scenario combinations. Relative changes were calculated to quantify the magnitude and direction of productivity changes over the simulation period. For each variable, the relative change between a given decade and the baseline period was expressed as a percentage:

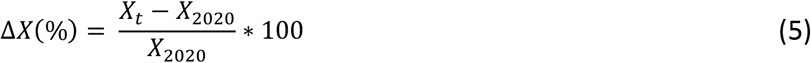

where *X_t_* represents the value in the target decade and *X_2020_* the corresponding value during the baseline decade (2015-2024). The same approach was applied to CV to quantify changes in interannual GPP variability relative to the baseline.

## 3. Results

### 3.1. Model evaluation

#### 3.1.1. Evaluation against remote-sensing GPP (2005-2014)

The ability of 3D-CMCC-FEM to reproduce historical GPP was evaluated against two independent remote-sensing products, GOSIF and MODIS, over the common period 2005-2014 (Vangi et al., 2025b). Across all Italian ecoregions, annual GPP generally ranged between approximately 500 and 2,000 gC m⁻² yr⁻¹, and the model reproduced the overall magnitude and spatial variability of forest productivity. Simulated GPP generally fell within the range of the remote-sensing estimates, although values were lower than both GOSIF and MODIS in most ecoregions, whereas a slight overestimation was observed in the Alpine region. GOSIF generally showed higher GPP values than MODIS, while the three climate forcings produced broadly comparable distributions. Among them, ERA5-Land generally produced median GPP values closest to the remote-sensing products (Fig. S1). The temporal evolution of simulated GPP was broadly consistent with both remote-sensing datasets, although discrepancies were more evident during the early part of the historical period and decreased towards the end of the time series (Fig. S2). The probability density functions (PDF) further showed substantial overlap between simulated and remote-sensing GPP distributions, despite differences in their central values and spread (Fig. S3).

ERA5-Land consistently showed the highest agreement with both remote-sensing products across the five ecoregions. Correlations (R) with GOSIF ranged from 0.68 to 0.96, however lower correlations were obtained against MODIS (R = 0.33-0.70). For ERA5-Land, the highest agreement was obtained in the Tyrrhenian ecoregion, while the lowest correlations occurred in the Alpine region. The model generally underestimated GPP in the Adriatic, Apennine, Po Plain, and Tyrrhenian ecoregions, with the largest negative biases occurring in the Apennine against GOSIF and in the Adriatic against MODIS. GFDL-ESM4 generally resulted in lower correlations and larger errors than ERA5-Land, while UKESM1-0-LL showed intermediate performance. Further details on the GPP model evaluation are provided in the Supplementary Material (Section 2.1.1, Evaluation against remote-sensing GPP).

#### 3.1.2. Evaluation against Italian NFI 2015

The model was further evaluated against the 2015 INFI (INFC, 2021) to assess its capability to reproduce GSV, AGB, and CS, under the three climate forcings (ERA5-Land, GFDL-ESM4, and UKESM1-0-LL), and across the five Italian ecoregions. The PDF showed substantial agreement between simulations and observations, reproducing the central tendency and overall shape of the observed distributions (Fig. S4). ERA5-Land generally produced the closest agreement with INFI observations, whereas GFDL-ESM4 and UKESM1-0-LL yielded highly similar distributions and mean values. The comparison of ecoregion-averaged values further confirmed the consistency between simulated and observed structural attributes (Fig. S5). Taylor diagrams indicated moderate to strong correlations between simulated and observed spatial patterns, with R ranging from 0.52-0.86 for GSV, 0.54-0.83 for AGB, and 0.52-0.81 for CS. ERA5-Land consistently showed the highest agreement across the three structural variables (Fig. S6). The model generally underestimated GSV and AGB, whereas CS was predominantly overestimated. Overall, model performance was highest for GSV, followed by AGB and CS. Differences between GFDL-ESM4 and UKESM1-0-LL were generally small, indicating that the choice of climate forcing had a limited influence on the representation of present-day forest structural attributes.

The detailed results of the INFI-based model evaluation are reported in the Supplementary Material (Section 2.1.2, Evaluation against Italian NFI 2015).

### 3.2. Climate change projections

#### 3.2.1. Temporal patterns of carbon dynamics across Italian ecoregions (2015-2100)

Future climate projections revealed distinct long-term responses of GPP flux and carbon stocks across the five ecoregions, with both the climate change scenarios and the driving ESMs shaping the projected trajectories (Fig. 3). During the first decades of the simulation (2020-2050), GPP, AGB, GSV, and CS followed broadly similar trajectories under the three scenarios: SSP126, SSP370, and SSP585. Thereafter, responses progressively diverged, with the separation among climate change scenarios becoming increasingly pronounced towards the end of the century (2050-2100). Although the simulations driven by GFDL-ESM4 and UKESM1-0-LL showed consistent overall temporal patterns, noticeable differences emerged in the magnitude of projected changes across several ecoregions, particularly under the highest-emission SSP585 scenario, highlighting the sensitivity of projected forest responses to the choice of climate forcing used to drive the 3D-CMCC-FEM simulations.

**Figure 3:**
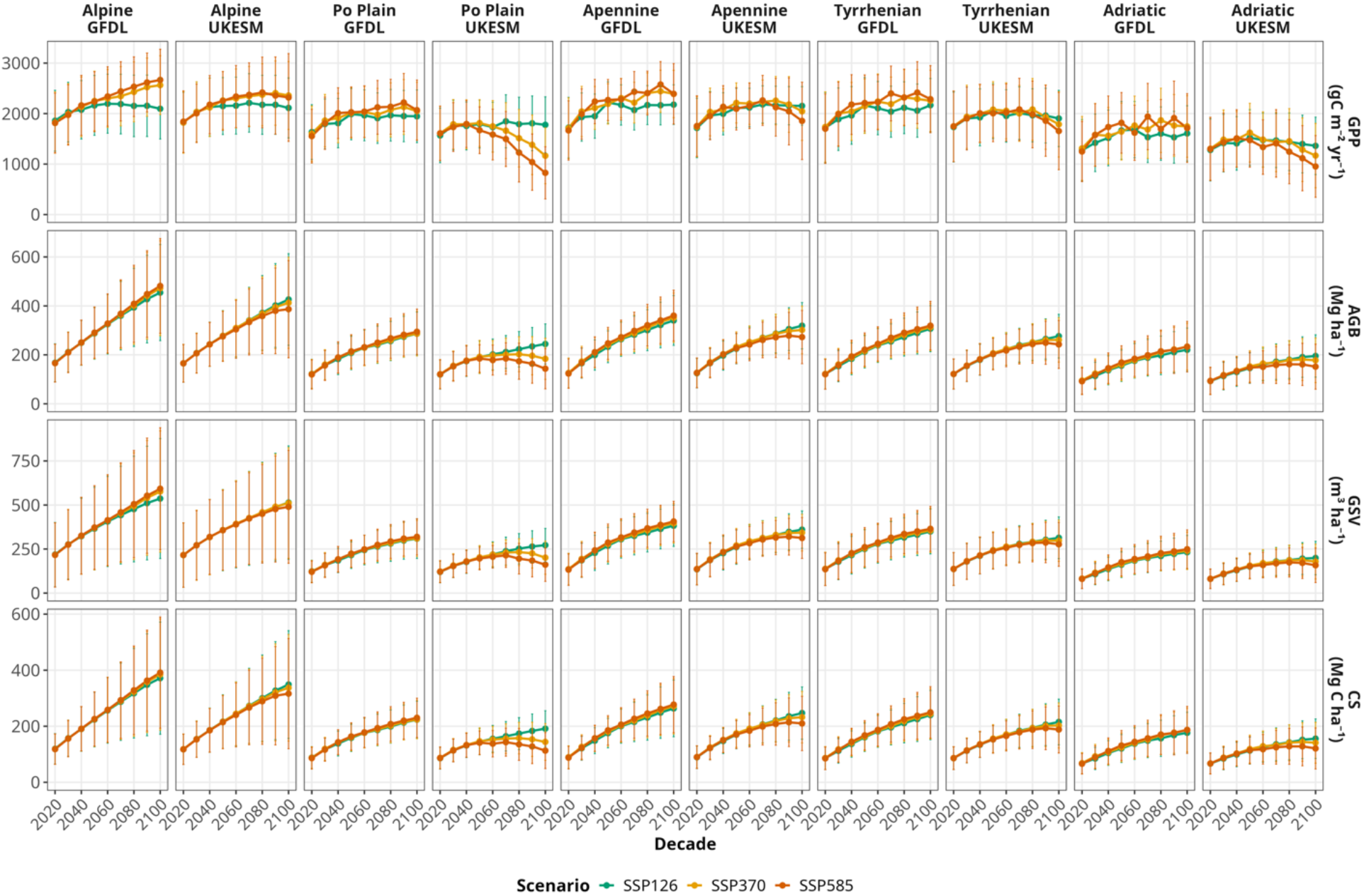
Decadal changes in forest productivity and carbon stocks. Mean decadal values (±1 standard deviation) of GPP (gC m⁻² yr⁻¹), AGB (Mg ha⁻^1^), GSV (m⁻^3^ ha⁻^1^), and CS (MgC ha⁻^1^) simulated from 2020 to 2100 for the five Italian ecoregions (i.e., Alpine, Apennine, Adriatic, Po Plain, and Tyrrhenian) under SSP126, SSP370, and SSP585. Results are presented separately for the GFDL and UKESM climate forcings.

GPP exhibited the strongest and earliest response to future climate change (Fig. 3). Under SSP126, GPP remained relatively stable throughout the century in all ecoregions, with only modest increases in the Alpine and Apennine regions and slight fluctuations elsewhere (between 1,500-2,500 gC m⁻² yr⁻¹). Considering the scenario SSP370, productivity generally increased during the first half of the century before stabilizing or slightly declining after approximately 2060-2070. In contrast, SSP585 produced clearly contrasting responses among ecoregions. Alpine forests showed a continuous increase in GPP, rising from approximately 1,800 to more than 2,500 gC m⁻² yr⁻¹ by the end of the century. Apennine and Tyrrhenian forests also experienced moderate increases during the first decades, reaching peak values around the 2070s before gradually declining towards 2100. By contrast, the Po Plain exhibited the strongest reduction in productivity, particularly under the UKESM1-0-LL forcing, where GPP declined from approximately 1,800 to less than 1,000 gC m⁻² yr⁻¹ by 2100. Adriatic forests displayed a similar decline, although less pronounced, decreasing from approximately 1,500 to around 1,100 gC m⁻² yr⁻¹ under SSP585. Despite these differences among climate scenarios and ecoregions, both ESMs reproduced the same qualitative responses, although differences in the magnitude of projected GPP became increasingly evident under SSP585. Compared with UKESM1-0-LL, GFDL-ESM4 generally projected higher and more variable productivity peaks during the first half of the century, reaching approximately 2,500 gC m⁻² yr⁻¹ under SSP585, whereas UKESM1-0-LL exhibited a more pronounced and persistent decline during the second half of the century, particularly in the Po Plain and Adriatic ecoregions.

In contrast to GPP, biomass and carbon stock variables responded more gradually to future climate change. CS has a remarkably similar temporal trajectory to AGB and GSV, characterized by a progressive increase throughout most of the simulation period under all climate scenarios (Fig. 3). Alpine forests consistently accumulated the largest biomass and carbon stocks, with AGB increasing from 200 to nearly 500 Mg ha⁻¹, GSV from 250 to more than 500 m³ ha⁻¹, and CS from 150 to nearly 400 MgC ha⁻¹ by the end of the century. Apennine and Tyrrhenian forests showed intermediate accumulation rates, reaching about 400 Mg ha⁻¹ for AGB (GSV: 500 m³ ha⁻¹; CS: 250-300 MgC ha⁻¹), whereas the Adriatic and Po Plain ecoregions maintained comparatively lower stock values, generally remaining below 300 Mg ha⁻¹ for AGB, 400 m³ ha⁻¹ for GSV, and 250 MgC ha⁻¹ for CS throughout the century. Differences among climate scenarios became increasingly evident after the mid-century. Under SSP126, biomass and carbon stocks increased steadily across all ecoregions, whereas SSP370 showed a progressive reduction in accumulation rates. The strongest response occurred under SSP585, where biomass accumulation slowed considerably, particularly in the Po Plain and Adriatic ecoregions, approaching a plateau or showing slight declines towards the end of the century. In contrast, Alpine and Apennine forests continued to accumulate biomass and carbon under all climate scenarios, indicating a sustained capacity for carbon sequestration despite changing climatic conditions. The simulations driven by GFDL-ESM4 and UKESM1-0-LL reproduced the same overall temporal patterns, although GFDL-ESM4 generally projected greater biomass accumulation, whereas UKESM1-0-LL simulated a more pronounced slowdown during the second half of the century under SSP370 and SSP585, particularly in the Po Plain and Adriatic ecoregions.

While reductions in GPP became evident after mid-century under the more severe scenarios, especially in the lowland and coastal ecoregions, biomass and carbon stocks generally continued to increase. The relative changes reported in Fig. S7 confirm the temporal patterns described above by quantifying the magnitude of the projected responses relative to the 2020 baseline. They highlight the progressive divergence among climate scenarios after mid-century and the different patterns of the five Italian ecoregions towards the end of the century. In addition, the decadal evolution of Current Annual Increment (CAI) and Mean Annual Increment (MAI) presented in Fig. S8 further supports these patterns, showing a progressive decline in stand productivity across all ecoregions throughout the century. Differences among climate scenarios became increasingly evident after mid-century, with the steepest reductions occurring under SSP585, particularly in the Po Plain and Adriatic ecoregions, whereas Alpine forests maintained the highest CAI/MAI values throughout the simulation period.

#### 3.2.2. Temporal patterns of carbon dynamics across forest types (2015-2100)

Projected responses to future climate change differed distinctly among forest types and largely reflected their distribution across the five ecoregions. Accordingly, the responses of individual forest types closely reflected the contrasting productivity trajectories previously identified across these ecoregions (Fig. 4). ‘*Mixed’* forests, which represented the largest proportion of the simulated stands and occurred across all ecoregions, maintained positive productivity trends under all climate scenarios. However, the magnitude of the response differed between the two climate forcings, with median GPP increasing by around 200-700 gC m⁻² yr⁻¹ under GFDL-ESM4, compared to 70-200 gC m⁻² yr⁻¹ under UKESM1-0-LL from 2020 to 2100. Forest types associated with the Alpine and Apennine ecoregions generally exhibited the greatest productivity throughout the century. Among broadleaved species, *F. sylvatica* and *Q. cerris* consistently increased under both ESMs, although GFDL-ESM4 projected substantially larger gains under SSP370 and SSP585. Under SSP585, median GPP increased by around of 800 and 1,200 gC m⁻² yr⁻¹ for *F. sylvatica* nd *Q. cerris*, respectively, under GFDL-ESM4, compared with 500 and 300 gC m⁻² yr⁻¹ under UKESM1-0-LL. Similarly, Alpine conifers responded positively to future climate change. *L. decidua* exhibited the strongest increase among all forest types, with GPP increasing by round 1,100-1,400 gC m⁻² yr⁻¹ under GFDL-ESM4 and UKESM1-0-LL, respectively, and considering the SSP585 climate scenario. *P. sylvestris*, *A. alba*, and *P. abies* also maintained sustained productivity increases, particularly under SSP370 and SSP585, although increases projected by UKESM1-0-LL were generally smaller than those simulated by GFDL-ESM4. In opposite, forest types of characteristics of the Tyrrhenian, Adriatic, and Po Plain ecoregions exhibited a much stronger dependence on the forcing climate model. With GFDL-ESM4, and SSP585, *Q. ilex* and *P. halepensis* showed positive productivity trends across all scenarios, with GPP increasing by up to around 500 and 600 gC m⁻² yr⁻¹, respectively. Conversely, UKESM1-0-LL projected progressive declines for both species, reaching −450 and −350 gC m⁻² yr⁻¹ by 2100. Likewise, *C. sativa* and *P. nigra* exhibited declining productivity under the higher-emission scenarios, with reductions consistently greater under UKESM1-0-LL than under GFDL-ESM4.

**Figure 4:**
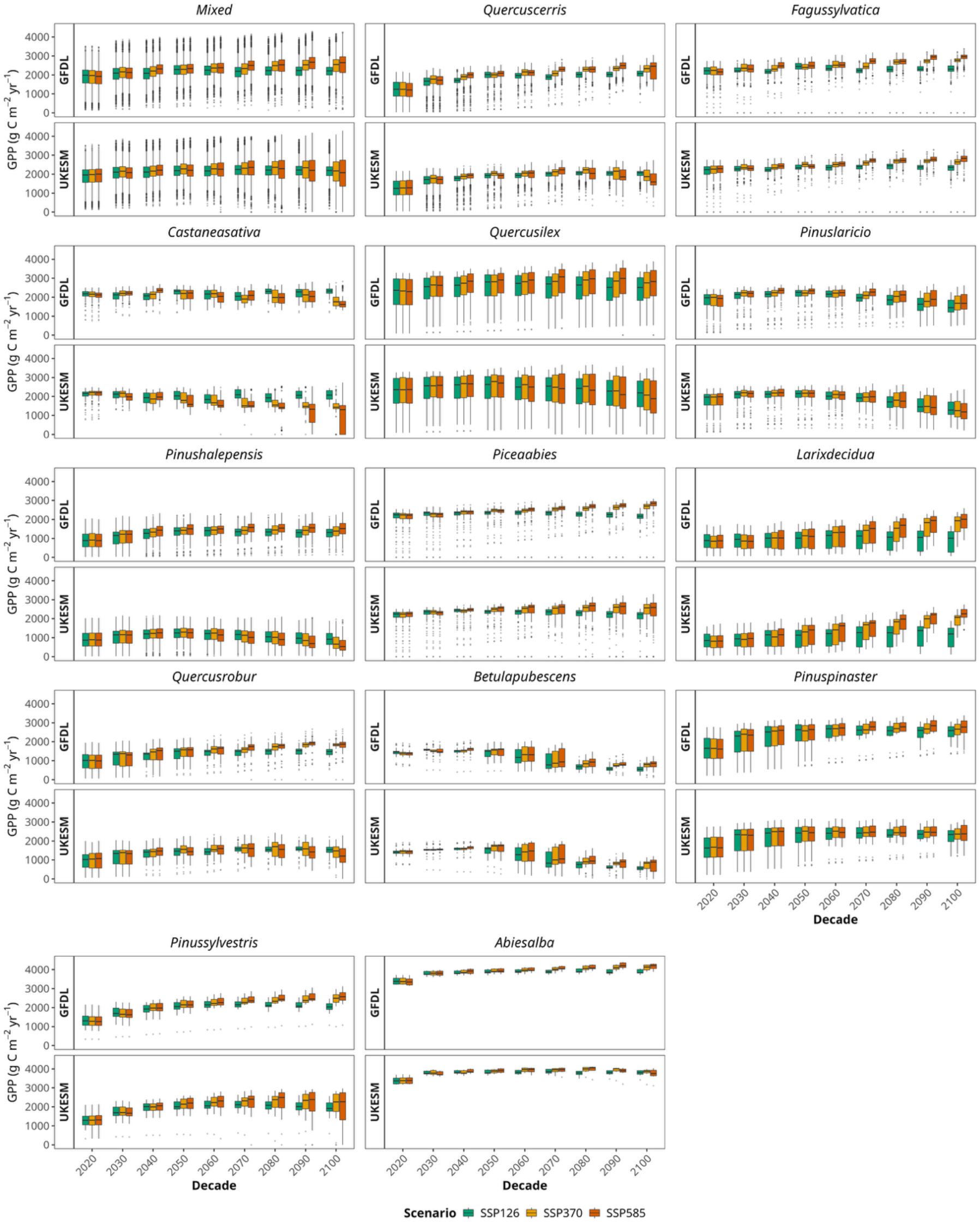
Decadal changes in GPP across the dominant Italian forest types under 3 climate change scenarios (SSP126, SSP370, and SSP585) during the period 2020-2100. Results are presented separately for the GFDL and UKESM climate forcings. Boxplots summarize the distribution of simulated GPP values (gC m⁻² yr⁻¹) across all NFI stands belonging to each forest type for each decade. The central line represents the median, the box the interquartile range (IQR), whiskers extend to 1.5 × IQR, and points denote outliers.

The temporal evolution of carbon stocks (Fig. S9) broadly mirrored the productivity responses (Fig. 4). Forest types of dominant in the Alpine and Apennine ecoregions, together with *‘Mixed’* forests, consistently accumulated the largest carbon stocks under both climate forcings throughout the simulation period. In contrast, forest types associated with the Tyrrhenian, Adriatic, and Po Plain ecoregions maintained comparatively lower carbon stocks, with smaller increases over time, particularly under SSP585. Differences between the two climate forcings were generally less pronounced for carbon stocks than for GPP, although UKESM1-0-LL projected lower end-of-century values for several coastal and lowland forest types.

#### 3.2.3. Temporal variability and stability of forest GPP across Italian ecoregions (2015-2100)

Figure 5 presents the temporal evolution of the CV of GPP, highlighting differences in the temporal variability and associated stability of forest GPP among climate scenarios, ecoregions, and forcing driven ESMs. Across all ecoregions, CV generally declined during the first half of the century (2020-2050), indicating an overall increase in GPP stability during this period, although the magnitude of this decrease varied according to the climate forcing and emission scenario. Differences among scenarios became progressively more evident after mid-century (2050-2100), particularly under SSP585, when several ecoregions exhibited increasing temporal variability in GPP and therefore declining ecosystem carbon flux stability. In contrast, SSP126 generally maintained CV values close to or below those of the baseline period (2015-2025), indicating relatively stable or increasing GPP stability, whereas SSP370 showed an intermediate response, with CV remaining rather steady until approximately 2050 before progressively diverging during the second half of the century.

**Figure 5:**
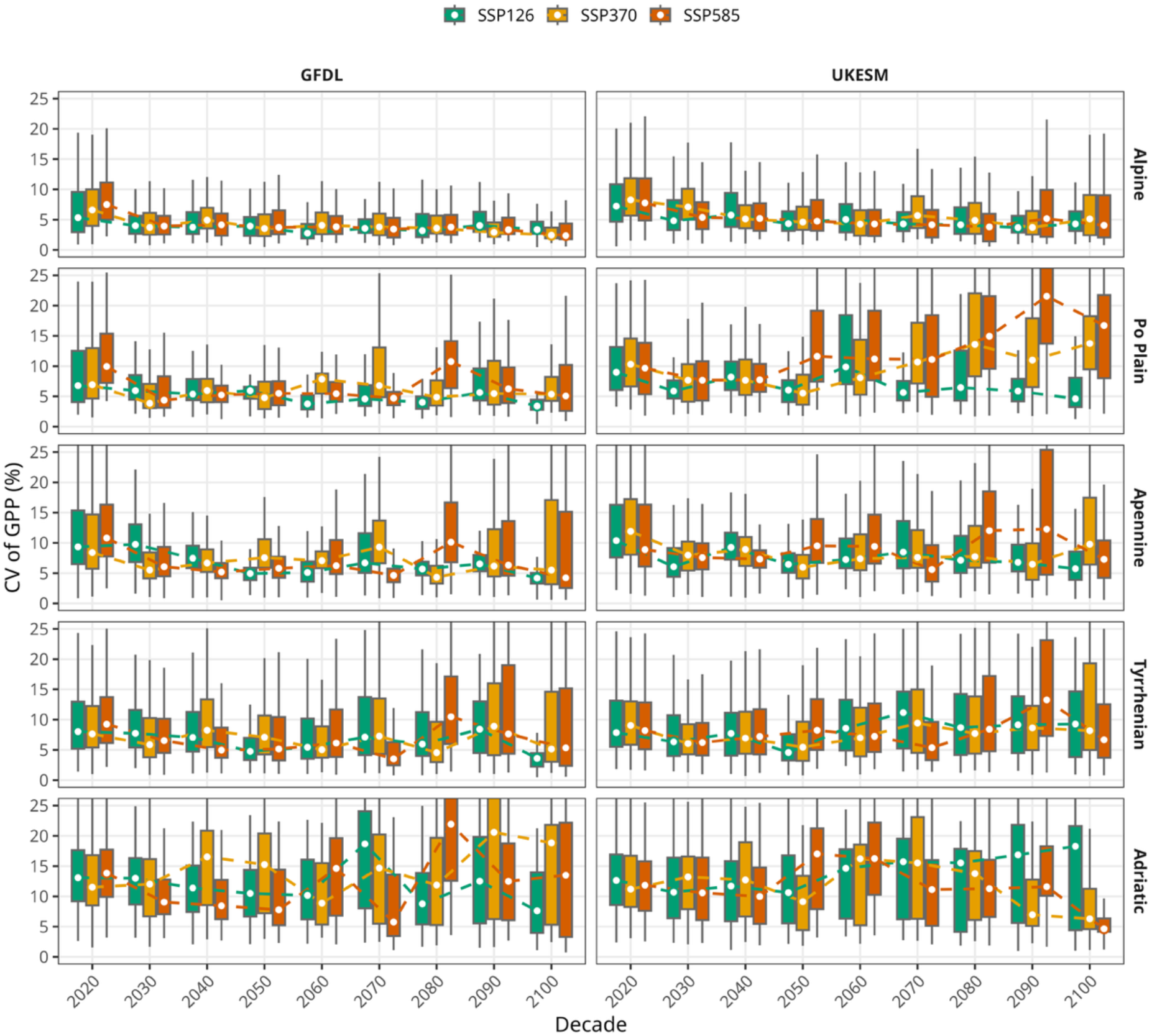
Decadal distributions of the coefficient of variation (CV, %) of GPP across forest stands within each Italian ecoregions (i.e., Alpine, Apennine, Adriatic, Po Plain, and Tyrrhenian) under SSP126, SSP370, and SSP585 scenarios. Boxplots represent the spatial variability among simulated stands, while dashed lines connect the decadal means. Results are shown separately for GFDL-ESM4 and UKESM1-0-LL climate forcings.

Noticeable differences emerged among ecoregions (Fig. 5; Table S7). Alpine forests consistently exhibited the lowest temporal variability and therefore the highest relative GPP stability under both climate forcings and all climate scenarios. Considering the GFDL-ESM4 climate forcing, mean CV decreased from approximately 7-9% in 2020 to 3-4% by 2100, corresponding to relative reductions of around 50%. UKESM1-0-LL projected a smaller decrease, with ΔCV ranging from −6 to −35%, indicating that Alpine forests maintained comparatively stable GPP flux even under SSP585. An intermediate pattern was observed in the Apennine ecoregion, with CV generally decreasing from 11-13% at the beginning of the century to 7-12% by 2100, irrespective of the climate forcing. This overall decline in CV indicates increasing temporal stability of GPP, with both ESM-driven simulations projected an overall reduction in temporal variability, with the strongest decline occurring under SSP126. Under this scenario, ΔCV reached −62% and −42% for GFDL-ESM4 and UKESM1-0-LL, respectively. A similar response was observed in the Tyrrhenian ecoregion, where CV remained relatively stable throughout the simulation period. This implies relatively constant stability of GPP over the century with GFDL-ESM4 forcing producing a consistent decline in CV across all climate scenarios (relative reductions ranging from −9 to −61%), whereas UKESM1-0-LL projected values close to the baseline, with slight increases under SSP370 (14%) and SSP585 (+8%). The latter increases indicate a modest decline in GPP stability under these scenarios. By contrast, Adriatic forests showed distinct responses depending on the driving ESM. Simulations forced by GFDL-ESM4 maintained relatively stable CV values throughout the century, generally fluctuating between 12 and 15%, with end-of-century changes ranging from −34% under SSP126 to 15% under SSP370, indicating moderately changing GPP stability depending on the scenario. In contrast, UKESM1-0-LL produced substantially greater temporal variability, characterized by wider boxplot distributions after mid-century, resulting in greater uncertainty and generally lower GPP stability, although end-of-century responses differed among scenarios, ranging from −33% (SSP585) to 6% (SSP126). The strongest response was observed in the Po Plain, where the two climate forcings projected markedly contrasting trajectories in GPP stability. Under GFDL-ESM4 forcing data, CV progressively declined throughout the century under all emission scenarios, with relative reductions ranging from 18 to 58% (increase in GPP stability). On the other hand, UKESM1-0-LL projected a pronounced increase in productivity variability under the higher-emission pathways (decline in GPP stability). Under SSP370, CV increased from 11.1% to 19.3% by the end of the century, corresponding to a 74% increase relative to the baseline period. An even stronger response occurred under SSP585, where CV rose from 10.2% to 23.6%, representing a 132% increase and indicating the strongest projected reduction in GPP stability among the ecoregions considered.

The relative changes in CV presented in Fig. S10 further quantified these temporal patterns relative to the baseline period, showing that ΔCV remained close to zero or negative under SSP126 (increasing GPP stability), whereas progressively larger positive changes emerged after approximately 2050 under SSP370 and particularly SSP585 (declining GPP stability). These changes were most pronounced in the Po Plain and, to a lesser extent, in the Tyrrhenian ecoregion under the UKESM1-0-LL forcing. The spatial distribution of ΔCV shown in Fig. S11 was consistent with these temporal patterns, revealing that increases in GPP variability and associated reductions in GPP stability were concentrated in the Po Plain and portions of the Tyrrhenian ecoregion under the higher-emission scenarios, while Alpine forests consistently exhibited limited changes or localized reductions in GPP variability and therefore maintained comparatively high GPP stability.

## 4. Discussion

### 4.1. Model Evaluation

#### 4.1.1. Evaluation against remote-sensing based GPP (2005-2014)

Simulations driven by ERA5-Land consistently provided the strongest agreement between simulated and remote sensing-derived GPP across the Italian ecoregions, outperforming those driven by GFDL-ESM4 and UKESM1-0-LL. This finding is consistent with previous evidence that the selection of meteorological forcing can considerably affect the performance of process-based simulations of terrestrial carbon fluxes, including those obtained with 3D-CMCC-FEM (Collalti et al., 2018; Dalmonech et al., 2024, 2026; Vangi et al., 2025). More broadly, comparisons of land-surface and carbon-cycle simulations have shown that atmospheric forcing accounted for approximately 75% of the variability in simulated global GPP (Hardouin et al., 2022), while studies comparing reanalysis products have reported improved land-surface performance when using ERA5 relative to earlier reanalysis (Albergel et al., 2018; Hu et al., 2022; Muñoz-Sabater et al., 2021). The stronger performance of ERA5-Land may partly reflect its finer spatial resolution (∼9 km) compared with the other forcings (∼50 km), which better captures Italy’s strong topographic and climatic gradients. These differences in the representation of temperature, radiation, and water availability can propagate into simulated GPP (Hardouin et al., 2022; Malle et al., 2025). Overall, these results highlight the role of climate forcing in shaping simulated GPP. The close agreement obtained under ERA5-Land provides an assessment of the model’s capacity to reproduce GPP under an observed-based climate forcing. In contrast, the differences in model performance under GFDL-ESM4 and UKESM1-0-LL during the historical period provide important context for interpreting their future projections, as differences in the representation of historical climate conditions may propagate into simulated GPP trajectories. The use of two alternative ESM forcings, therefore, provides a representation of forcing-related uncertainty in future GPP projections and captures a broader range of potential GPP responses.

#### 4.1.2. Evaluation against Italian NFI 2015

The 3D-CMCC-FEM closely reproduced the probability distributions, means and standard deviations of GSV, AGB and CS observed in the 2015 INFI (INFC, 2021), capturing their central tendency, spread and spatial variability across Italy. ERA5-Land showed the closest agreement, while GFDL-ESM4 and UKESM1-0-LL produced highly similar results, indicating a consistent representation of present-day forest structure across different climate forcings. These results are consistent with the national-scale validation conducted by Vangi et al.(2025b) against the third Italian NFI, which demonstrated good performance in reproducing forest structural attributes and carbon stocks across Italy. The present analysis extends this evidence by demonstrating that agreement with observed forest structure is maintained under the alternative climate forcings adopted here. In addition, initialization from observed forest structure constrains the simulated starting conditions and helps preserve spatial heterogeneity. Overall, the close agreement with the INFI provides a robust present-day structural and carbon-stock baseline for assessing future changes in forest biomass and carbon stocks.

### 4.2. Climate change projections

#### 4.2.1. Temporal patterns of carbon dynamics across Italian ecoregions (2015-2100)

Carbon flux followed broadly similar trajectories across SSP126, SSP370 and SSP585 until approximately mid-century, after which scenario differences became increasingly pronounced, particularly under SSP585. This temporal divergence is consistent with previous projections at both European and Italian scales, which indicate that GPP trajectories remain relatively similar during the first half of the century but increasingly diverge thereafter, with higher-emission scenarios generally maintaining higher productivity while SSP126 progressively levels off (Jones et al., 2023; Knauer et al., 2023; Zhou et al., 2024). For Italy, however, increasing warming and reduced precipitation are expected to progressively constrain GPP during the second half of the century as the positive effects of atmospheric CO₂ fertilization weaken under increasing temperature and water limitations (Fibbi et al., 2019; Testolin et al., 2023). The divergence among scenarios therefore likely reflects the cumulative effects of increasingly contrasting thermal and water conditions, which may eventually lead to severe productivity declines, carbon starvation and mortality in vulnerable stands under extreme climatic stress (Fibbi et al., 2019; Saponaro et al., 2025; Sazeides and Fyllas, 2025).

Carbon stocks showed broadly similar trajectories across scenarios until approximately 2050, followed by a more gradual divergence (Karjalainen et al., 2003; Luo et al., 2025; Pilli et al., 2026, 2022; Qiu et al., 2025). Compared with GPP, differences among scenarios remained smaller, reflecting the slower response of carbon stocks, which integrate changes in productivity, respiration and carbon allocation over longer timescales (Bennett et al., 2024; Cao and Woodward, 1998; Sun et al., 2024). Consequently, forest carbon stocks can continue to increase even when carbon fluxes begin to diverge, although prolonged climatic stress may eventually lead to carbon losses when respiration exceeds carbon uptake (Collalti et al., 2018; Finzi et al., 2020; Yue Li et al., 2022). The temporal dynamics therefore suggest that changes in carbon fluxes may provide an earlier indication of emerging climate constraints than changes in forest carbon stocks.

The projected biomass and carbon-stock trajectories also showed strong heterogeneity among ecoregions. Alpine forests generally accumulated the largest biomass and carbon stocks, followed by Apennine and Tyrrhenian forests, whereas Adriatic and Po Plain forests maintained lower values. This pattern is broadly consistent with evidence of high carbon storage in Alpine forests, while also reflecting the importance of forest type, stand structure and age, which can generate substantial within-ecoregion variability (Bono et al., 2024; Gasparini and Di Cosmo, 2015; Keith et al., 2024; Vangi et al., 2024a). Under the more severe emission scenarios (i.e., SSP370 and SSP585), Alpine and Apennine forests maintained positive biomass and carbon accumulation, whereas accumulation slowed significantly in the Adriatic and Po Plain regions. The finding is coherent with previous studies showing that mountain forests can maintain relatively high carbon accumulation under severe climate change (Blattert et al., 2024; Oddi et al., 2022; Walter et al., 2025). In particular, evidence from the Apennines suggests that species complementarity can contribute to maintaining carbon accumulation under SSP370, while simulations for the Italian Alps indicate relatively limited differences in carbon stocks between RCP4.5 and RCP8.5 (Blattert et al., 2024; Brocco et al., 2026; Walter et al., 2025). By contrast, warmer lowland and Mediterranean forests are generally more exposed to increasing heat and water stress, which can progressively constrain productivity and carbon accumulation (Gea-Izquierdo et al., 2017; Lindner et al., 2010).

The magnitude of projected changes in carbon stocks and productivity under the different emission scenarios is likely influenced not only by elevation, forest composition, and local environmental conditions, but also by climate forcing. GFDL-ESM4 and UKESM1-0-LL generally reproduced the same qualitative temporal patterns, but increasingly diverged in the magnitude of projected changes, particularly under SSP585, for which UKESM1-0-LL projected the strongest declines in carbon stocks and productivity. It is characterized by relatively high transient and effective climate sensitivity, resulting in stronger warming under high-emission scenarios and an earlier exceedance of warming thresholds compared with many CMIP6 models (Mulcahy et al., 2023; Swaminathan et al., 2022). This provides a plausible mechanistic explanation for the stronger declines projected under UKESM1-0-LL, particularly under SSP585, where differences in the magnitude of climate forcing become increasingly pronounced (Tebaldi et al., 2021).

#### 4.2.2. Temporal patterns of carbon dynamics across forest types (2015-2100)

Projected GPP responses differed among forest types and broadly reflected the contrasting carbon productivity trajectories observed across the Italian ecoregions. *‘Mixed’* forests and forest types associated with Alpine and Apennine environments generally maintain positive productivity trends, including Alpine conifers, and Apennine broadleaved forests. In contrast, forest types of characteristics of the Tyrrhenian and Adriatic ecoregions, mainly represented by coastal species, together with lowland forests of the Po Plain, generally showed a stronger dependence on the forcing ESM (See paragraph 2.1., Italian case study for the distribution of forest types and species).

This geographical distinction is already well documented in previous studies of European forests, which generally identify a north-south gradient in productivity responses, with mountain and northern forests benefiting from warming conditions, although warmer southern forests become increasingly constrained by climatic stress (Fibbi et al., 2019; Reyer et al., 2014; Trotsiuk et al., 2020). In particular, Fibbi et al., (2019) reported the strongest GPP increases for Alpine Mountain conifers, with increases approaching 40% under RCP4.5, while Reyer et al. (2014) projected *P. abies* and *P. sylvestris* L. to maintain positive productivity responses across central and northern Europe. Similarly, Trotsiuk et al., (2020) found that these two species in Switzerland increased productivity at higher elevations during warm-dry extremes. Among the Alpine conifers, the particularly strong response projected for *L. decidua*, under SSP585 is especially noteworthy, as recent evidence identifies *Larix*-dominated treelined systems among those showing the strongest increases in GPP under warming (Mallen-Cooper et al., 2026). The positive trajectories projected here for Alpine conifers and Apennine broadleaved species are therefore broadly consistent with ecoregion patterns. Again, such responses may reflect the partial alleviation of temperature limitation in colder environments, particularly at higher elevations, where warming can increase photosynthetic activity and extend the growing season (Blattert et al., 2024; Oddi et al., 2022; Walter et al., 2025). In addition to climatic gradients, the positive productivity trajectories projected for ‘*Mixed’* forests may also be consistent with the productivity advantage associated with species mixtures. A global meta-analysis of 126 case studies found that mixed-species stands were, on average, 15% more productive than the mean of their component mono-species (Jactel et al., 2018).

In contrast, forest types typical of the Tyrrhenian and Adriatic ecoregions, together with lowland forests of the Po Plain, showed a stronger dependence on the forcing ESMs. This dependence was particularly evident for Mediterranean coastal species, for which GFDL-ESM4 generally projected positive productivity trajectories, whereas UKESM1-0-LL resulted in substantially stronger declines. The contrasting responses of *Q. ilex* and *P. halepensis* between the two forcing ESMs suggest that productivity in these warmer environments is particularly sensitive to the climatic conditions imposed by the forcing model. Previous studies provide further evidence of this climate-sensitivity, showing that coastal forests become increasingly vulnerable as warming intensifies, particularly when higher temperatures are accompanied by reduced water availability (Dalmonech et al., 2026; Helman et al., 2017; Sazeides and Fyllas, 2025). For example, Helman et al. (2017) showed *P. halepensis* forests across the Mediterranean basin have been projected to experience GPP declines of approximately 16% and 31% under warming of 1°C and 2°C, respectively, independently of water-use strategy. This pattern is further illustrated by the response of *P. nigra*. The evident decline projected under UKESM1-0-LL, particularly towards the end of the century under SSP585, is consistent with the stronger warming and changes in precipitation projected by this forcing relative to GFDL-ESM4. These climatic differences become increasingly important under stronger warming, potentially pushing already warm and water-limited coastal environments beyond the conditions favorable for sustained productivity. The stronger response of coastal forests compared with their mountain counterparts further suggests that the climatic conditions imposed by the forcing ESM can be especially consequential for forest types already close to their thermal limits (Mulcahy et al., 2023; Swaminathan et al., 2022).

The temporal dynamics of carbon stocks generally followed the patterns observed of GPP, with forest types showing increasing productivity generally also accumulating carbon over time. However, carbon stocks responded more gradually, reflecting their integration of productivity, respiration, and carbon allocation over longer timescales (Bennett et al., 2024; Cao and Woodward, 1998; Sun et al., 2024). Consequently, differences among climate forcings and scenarios were less pronounced for carbon stocks than for GPP.

#### 4.2.3. Temporal variability and stability of forest GPP across Italian ecoregions (2015-2100)

The coefficient of variation of annual GPP was used to quantify variability in ecosystem productivity. The CV is widely used to quantify relative temporal variability as it normalizes year-to-year fluctuations by background productivity and makes comparisons across sites and forest types more interpretable. This use is scientifically well grounded, but also partial. By scaling variability by mean productivity, CV provides a useful, normalized metric for comparing ecosystems with significantly different baseline productivity (Li et al., 2024; Lin et al., 2023; Zhang et al., 2022). However, CV provides an indicator of temporal stability based on the magnitude of variability, rather than a mechanistic explanation of the processes underlying stability. Across forest ecosystems, variability in GPP reflects the combined effects of climate forcing, phenology, structural attributes, and modelling design. Therefore, CV summarizes the magnitude of variability without identifying its specific causes (De Pue et al., 2023; Xia et al., 2015; Yao et al., 2018). Accordingly, several studies complement CV with trend and anomaly analyses or phenological metrics (De Pue et al., 2023; Yuzhen Li et al., 2022; Yao et al., 2018), and overall, the literature indicates that forest GPP variability is climate-sensitive but not controlled by a single climatic driver (Yuzhen Li et al., 2022; Li et al., 2020; Yao et al., 2018).

The results revealed clear spatial differences in the temporal stability of GPP across Italian ecoregions, as indicated by contrasting trajectories in the coefficient of variation (CV). The Po Plain showed the strongest decline in GPP stability, with the largest increases in GPP variability under higher-emission scenarios (i.e., SSP370 and SSP585), particularly under UKESM1-0-LL, while Alpine forests consistently exhibited the highest temporal stability of GPP, with the lowest temporal variability across scenarios and climate forcings and CV generally declining throughout the century. Tyrrhenian and Apennine forests showed intermediate patterns, while Adriatic forests generally exhibited higher variability and consequently lower GPP stability. This regional differentiation is consistent with the contrasting climatic constrains on forest productivity across Italy. Fibbi et al. (2019) showed that forest growth in the driest peninsular areas is primarily constrained by water availability, whereas temperature is the dominant limitation at higher elevations. Alpine forests may therefore maintain relatively high GPP stability where warming could reduce temperature limitation without inducing strong water stress. In contrast, peninsular forests are more exposed to fluctuations in atmospheric water demand, which may contribute to greater GPP variability and lower temporal stability. Fibbi et al. (2019) further reported greater spatial heterogeneity in precipitation than temperature in the peninsular areas, with reductions particularly concentrated during spring and summer, while precipitation remained comparatively stable in the northern Alps.

The pronounced increase in CV observed in the Po Plain and the associated decline in GPP stability may therefore reflect, at least in part, differences in the temporal variability and intensity of the climatic conditions imposed by the different forcing. Although direct evidence for Po Plain GPP variability under SSP585 is limited, Mediterranean forest studies indicate that increasing temperature and vapor pressure deficit, together with reduced rainfall, can negatively affect GPP (Lombardi et al., 2023). The contrasting CV trajectories between GFDL-ESM4 and UKESM1-0-LL further suggest that differences in the temporal evolution of the climate forcing can propagate into ecosystem productivity variability and consequently into contrasting trajectories of GPP stability. The contrast among ecoregions may thus reflect both differences in the climatic variability imposed by the ESMs climate forcing and differences in how ecosystems respond to this variability, resulting in different levels and trajectories of GPP stability. This interpretation is also supported by global evidence showing greater GPP variability in water-limited ecosystems, where water availability represents a major control on productivity (O’Sullivan et al., 2020). The increasing influence of water availability on GPP at annual and longer timescales further supports the importance of hydrological constraints for future productivity variability and GPP stability (Liang et al., 2024). Moreover, terrestrial ecosystem models have shown to tend to underestimate GPP variability, particularly in water-limited ecosystems (Lin et al., 2023), suggesting that the projected increase in variability in lowland forests may be conservative and that the associated decline in GPP stability could be underestimated.

Climate extremes and episodic stress events may also contribute to changes in GPP stability. Climate extremes can substantially affect GPP by increasing variability in carbon uptake and reducing forest functional stability (O’Sullivan et al., 2020; Qin et al., 2026). In particular, compound hot-dry events have been identified as an important driver of negative GPP anomalies and increased carbon variability across forests, with such events accounting for more than 40% of negative GPP extremes in CMIP6 Earth system model projections. The effects of climate extremes also depend on forest composition, stand structure, and elevation, with different forest types showing contrasting resistance to climatic stress, potentially contributing to the contrasting patterns of GPP stability observed among Italian ecoregions (Gampe et al., 2021; Gao et al., 2024).

## 5. Limitations

The modelling framework presented here includes several modelling choices and limitations that should be acknowledged when interpreting the projected changes in forest carbon dynamics. Anthropogenic disturbances, including forest management and harvesting, were not explicitly simulated as a deliberate modelling choice, allowing the analysis to isolate the effects of climate change on forest ecosystems without introducing additional variability associated with future management decisions. Another limitation is the absence of major natural disturbances, such as fire, insect outbreaks, and diseases were not represented. These processes can substantially alter both carbon fluxes and stocks and may therefore modify the magnitude and spatial pattern of the projected climate-driven responses. The absence of disturbance processes may consequently lead to an overestimation of carbon accumulation where future disturbance frequency or severity increases (Grünig et al., 2026). Other environmental stressors were also not considered, including potential changes in nitrogen availability, ozone effects, and their interactions with rising atmospheric CO₂ concentration. Such processes may either amplify or partially offset the impact of climate change on forest productivity and carbon storage. Finally, species migration and range shifts were not explicitly simulated. Changes in species composition resulting from the dispersal, establishment, and expansion of species currently absent from a stand may occur over longer timescales than those represented here. Such changes could alter stand composition and potentially lead to a different species becoming dominant under future climatic conditions.

## 6. Conclusions

We applied a detailed biogeochemical, biophysical, process-based forest model to assess the potential impact of climate change on carbon gross productivity, carbon stocks, on the stability of Italian forests from 2005 to 2100 under the ISIMIP framework. Evaluation against independent remote sensing-based GPP ddata andforest inventory observations demonstrated the model’s capability to reproduce the main spatial and temporal patterns of both forest productivity and structural carbon pools, providing a consistent baseline for assessing future climate-driven changes.

Simulations under a portfolio of diverse climate change scenarios revealed pronounced differences among Italian ecoregions and forest types. Alpine and Apennine forests generally maintained positive productivity trajectories and continued to accumulate biomass and carbon under future climate scenarios, whereas Mediterranean coastal and lowland forests showed stronger constraints, particularly under higher-emission climate scenarios. Across forest types, mountain species (i.e., Alpine conifers and Apennine broadleaved forests) generally maintained stronger and more stable GPP trajectories and carbon accumulation, while lowland and coastal species exhibited greater variability and stronger declines, particularly under the warmer forcing, reflecting their greater exposure to climatic stress. This greater sensitivity may reflect not only their higher exposure to climatic stress, but also their proximity to climatic thresholds, such that relatively small differences among climate forcings can lead to substantially different productivity trajectories. ‘*Mixed’* forests also maintained positive productivity trajectories, consistent with evidence of enhanced productivity in species mixtures.

The temporal dynamics of carbon stocks broadly followed those of GPP but responded more gradually, indicating that changes in carbon fluxes can emerge before corresponding changes in forest carbon pools. At the same time, variability in GPP revealed contrasting patterns of ecosystem stability across forest types. Mountain forests generally maintained lower and more stable gross productivity variability, whereas lowland and coastal forests showed greater increases in GPP variability, particularly under higher-emission scenarios and the warmer UKESM1-0-LL forcing, indicating greater sensitivity of forest systems to climatic change. Carbon stocks, however, continued to accumulate in several of these forest types despite increasing productivity variability, highlighting that changes in ecosystem stability may emerge before corresponding declines in carbon storage.

Finally, the contrasting trajectories simulated by GFDL-ESM4 and UKESM1-0-LL highlight climate-forcing uncertainty as an important component of future projections. This uncertainty was particularly pronounced in warm and potentially water-limited forests, where differences in projected temperature and precipitation translated into substantially different productivity trajectories, indicating that the magnitude of future climate impacts depends strongly on the climatic forcing applied. Accounting for both ecoregional differences in climate sensitivity and uncertainty among climate forcings is therefore essential when assessing future forest resilience and carbon sequestration potential. The identified differences among mountain, lowland, and coastal forests can help identify systems more exposed to future climate stress and inform the prioritization of targeted adaptation strategies.

