## Supplemental Material for "Climate change threatens the productivity stability of Italian forests"

#### 1. Material and Methods

##### 1.1. The 3D-CMCC-FEM model

###### 1.1.1. Model description

Forest growth and carbon dynamics were simulated using the 3D-CMCC-FEM (version 5.7), a process-based forest ecosystem model that simulates the dynamics of carbon, nitrogen, and water cycles under changing environmental conditions, forest management, and climate scenarios (Collalti et al., 2024, 2018, 2016; Dalmonech et al., 2022; Vangi et al., 2025a). Gross primary production (GPP) is simulated using the biochemical photosynthesis model of Farquhar et al. (1980), parameterized following (Bernacchi et al., 2003, 2001), and coupled with a multilayer "big-leaf" approach based on the exponential light attenuation (Monsi-Saeki formulation), which distinguishes between sunlit and shaded foliage (Collalti et al., 2018; De Pury & Farquhar, 1997; Medlyn et al., 2002). The acclimation of leaf photosynthesis to temperature is represented following Kattge and Knorr (2007), while temperature effects on enzyme kinetics are described through an Arrhenius relationship similarly adjusted for thermal acclimation (Collalti et al., 2018). Autotrophic respiration ($R_a$) follows the "Growth-and-Maintenance-Respiration-Paradigm", in which respiration is partitioned into maintenance respiration ($R_m$), associated with the metabolic costs of maintaining existing tissues, and growth respiration (Rg), associated with the synthesis of new tissues (Collalti et al., 2020). Net Primary Production (NPP) is calculated as the difference between GPP and $R_a$. Carbon assimilated through photosynthesis is allocated among structural biomass pools (leaves, stems, branches, fruits, fine roots and coarse roots) and a non-structural carbon (NSC) pool containing starch and soluble sugars. Allocation follows species-specific phenological rules and differs between evergreen and deciduous species. The NSC pool buffers temporal imbalances between carbon supply and demand and can sustain respiration and tissue development during periods of negative carbon balance (Collalti et al., 2017, 2020; Merganičová et al., 2019). The model represents forest structural development through the dynamics of tree cohorts and biomass pools, accounting for stand growth, competition for light, self-thinning, age-related mortality, and carbon starvation. Forest management can additionally modify stand structure and biomass through operations such as thinning and final harvesting (Dalmonech et al., 2022; Testolin et al., 2023).

#### 1.2. Experimental design

##### 1.2.1. Model evaluation

Model performance was evaluated against independent observations of carbon fluxes and carbon stocks using satellite-derived GPP estimates and INFC 2015 (INFC, 2021), respectively. All evaluations were conducted consistently across the three historical climate forcings (i.e., ERA5-Land, GFDL-ESM4, and UKESM1-0-LL) and the five Italian ecoregions (i.e., Adriatic, Alpine, Apennine, Po Plain, and Tyrrhenian).

Two complementary satellite-based GPP products were used: GOSIF and MODIS. The GOSIF product combines solar-induced fluorescence (SIF) observations from the Orbiting Carbon Observatory-2 (OCO-2) with the enhanced vegetation index (EVI) from the Moderate Resolution Imaging Spectroradiometer (MODIS), together with meteorological variables from the MERRA-2 reanalysis (Li and Xiao, 2019a, 2019b). The final GPP product was obtained by relating OCO-2 SIF to flux-tower GPP, providing annual GPP estimates at a spatial resolution of 0.05° (~5 km). The MODIS satellite-based product provides terrestrial GPP values based on the radiation-use-efficiency approach. The final product is generated at an 8-day temporal and 500 m spatial resolution from Terra and Aqua observations (Running and Zhao, 2021). For comparison with the annual model outputs, the 8-day GPP estimates were aggregated to annual values. Only observations meeting the highest product-quality criteria were retained, following the quality-control specifications of the MODIS product (Vangi et al., 2025b). For both products, annual GPP estimates were matched to the corresponding simulated stands and compared with 3D-CMCC-FEM outputs for 2005-2014 period.

Following the same approach described by Vangi et al. (2025), model performance was also evaluated against independent observations of forest carbon stocks and structural attributes from the INFC 2015 (INFC, 2021). The modelled forest state corresponding to 2015 was compared with inventory observations for three variables: GSV, AGB, and CS, which provide complementary information on forest structural development and carbon storage. Simulated values were compared with the corresponding inventory observations at the stand level and subsequently aggregated by ecoregion.

Model-observation agreement was quantified using five complementary statistical metrics: bias, mean absolute error (MAE), root mean square error (RMSE), and Pearson's correlation coefficient (r). These metrics were applied consistently to both remote-sensing and inventory evaluations, providing a common framework for assessing model performance across carbon fluxes and carbon stocks

**Table S1:** Overview of the climate forcing datasets used to drive the 3D-CMCC-FEM simulations for the 5135 Italian National Forest Inventory (INFI) stands (INFC, 2021). Simulations were performed using two Earth System Model (ESM) (GFDL-ESM4 and UKESM1-0-LL) under historical conditions (2005-2014) and three future climate scenarios (SSP126, SSP370, and SSP585; 2015-2100). The ERA5-Land reanalysis dataset was used as the reference historical climate forcing for the period 2005-2014.

| Plots | ESM | Climate Forcing datasets | Time Resolution |
|---|---|---|---|
| **5135 Italian NFI stands** | GFDL-ESM4 | HIST+SSP126 | 2005-2100 |
| | | HIST+SSP370 | 2005-2100 |
| | | HIST+SSP585 | 2005-2100 |
| | UKESM1-0-LL | HIST+SSP126 | 2005-2100 |
| | | HIST+SSP370 | 2005-2100 |
| | | HIST+SSP585 | 2005-2100 |
| | ERA5-Land | HIST | 2005-2014 |

### 2. Results

#### 2.1. Model Evaluation

##### 2.1.1. Evaluation against remote-sensing GPP (2005-2014)

The ability of 3D-CMCC-FEM to reproduce historical gross primary productivity (GPP) was evaluated by comparing simulated annual GPP with two independent remote sensing products, GOSIF and MODIS, over the common period 2005-2014 (Vangi et al., 2025b). Across all ecoregions and datasets, annual GPP generally ranged between approximately 500 and 2000 gC $m^{-2}$ $yr^{-1}$. The comparison of GPP distributions showed that the model successfully reproduced the overall magnitude and spatial variability of forest productivity across the five Italian ecoregions. Although differences in median GPP and distribution spread were evident among climate forcings and remote sensing products, simulated values generally fell within the observed range across all ecoregions (Fig. S1). Simulated GPP was generally lower than both GOSIF and MODIS in all ecoregions except the Alpine region, where a slight overestimation was observed. GOSIF consistently reported higher median GPP than MODIS, while the three climate forcings produced comparable distributions. Among them, ERA5-Land showed median values closest to the satellite products, whereas GFDL-ESM4 and UKESM1-0-LL exhibited a slightly larger spread, particularly in the Apennine, Po Plain, and Tyrrhenian ecoregions. Simulated and satellite-derived GPP exhibited similar temporal dynamics over the historical period, with larger discrepancies during the early years and progressively better agreement towards the end of the time series (Fig. S2). In addition, probability density functions (PDF) showed substantial overlap among datasets despite differences in their median values (Fig. S3).

Among the three climate forcings, ERA5-Land consistently provided the highest agreement with both remote sensing products across all ecoregions. Against GOSIF, correlation coefficients ranged from R=0.68-0.96, whereas lower correlations were obtained against MODIS (R=0.33-0.70). In both cases, model performance ranked as Tyrrhenian (GOSIF: R = 0.96; MODIS: R = 0.70) > Apennine (GOSIF: R = 0.89; MODIS: R = 0.63) > Adriatic (GOSIF: R=0.88; MODIS R=0.68) > Po Plain (GOSIF: R=0.76; MODIS: R=0.61) > Alpine (GOSIF R=0.68; MODIS: R=0.33). ERA5-Land systematically underestimated GPP in the Adriatic, Apennine, Po Plain, and Tyrrhenian ecoregions, with the largest negative bias occurring in the Apennine against GOSIF (−690.22 gC $m^{-2}$ $yr^{-1}$) and in the Adriatic against MODIS (−484.90 gC $m^{-2}$ $yr^{-1}$), whereas Alpine forests exhibited a slight positive bias for both products 86.71 and 241.88 gC $m^{-2}$ $yr^{-1}$, respectively. Prediction errors followed a similar pattern, with the highest MAE and RMSE observed in the Apennine against GOSIF (MAE=690.22; RMSE=707.61 gC $m^{-2}$ $yr^{-1}$) and in the Adriatic against MODIS (MAE = 484.90; RMSE = 504.15 gC $m^{-2}$ $yr^{-1}$). Conversely, Alpine forests showed the lowest prediction errors against GOSIF (MAE=135.76; RMSE=162.70 gC $m^{-2}$ $yr^{-1}$), while against MODIS the lowest MAE and RMSE were recorded in the Alpine ecoregion (MAE=245.83; RMSE=286.50 g C $m^{-2}$ $yr^{-1}$).

Simulations driven by GFDL-ESM4 reproduced the regional variability of GPP but showed consistently lower performance than ERA5-Land. Correlations with GOSIF ranged from R=0.40-0.88 and further decreased against MODIS (R=0.15-0.52). The Tyrrhenian ecoregion again showed the highest agreement, whereas Alpine forests exhibited the weakest correlations. Compared with ERA5-Land, GFDL-ESM4 generally increased the magnitude of the negative bias, reaching −629.22 gC $m^{-2}$ $yr^{-1}$ in the Apennine against GOSIF and −538.56 gC $m^{-2}$ $yr^{-1}$ in the Adriatic against MODIS, while the Alpine ecoregion remained the only region with a positive bias (68.94 and 224.11 gC $m^{-2}$ $yr^{-1}$ compared with GOSIF and MODIS, respectively). Prediction errors remained highest in the Apennine against GOSIF (MAE=629.22; RMSE=651.84 gC $m^{-2}$ $yr^{-1}$) and in the Adriatic against MODIS (MAE=538.56; RMSE=555.40 gC $m^{-2}$ $yr^{-1}$). Conversely, Alpine forests exhibited the lowest prediction errors for both products (GOSIF: MAE=161.27; RMSE=189.45 gC $m^{-2}$ $yr^{-1}$; MODIS: MAE=246.81; RMSE=297.89 gC $m^{-2}$ $yr^{-1}$).

UKESM1-0-LL showed intermediate performance between ERA5-Land and GFDL-ESM4. Correlations ranged from R=0.55-0.96 against GOSIF and R=0.14-0.59 against MODIS. Similar to the other climate forcings, the Tyrrhenian ecoregion consistently showed the highest agreement, whereas Alpine forests exhibited the weakest agreement. Biases and prediction errors followed spatial patterns comparable to those obtained with ERA5-Land and GFDL-ESM4. The largest negative bias was observed in the Apennine considering GOSIF (−619.95 gC $m^{-2}$ $yr^{-1}$) and in the Adriatic against MODIS products (−548.15 gC $m^{-2}$ $yr^{-1}$), whereas Alpine forests remained the only ecoregion with a positive bias (66.62 and 221.79 gC $m^{-2}$ $yr^{-1}$ against GOSIF and MODIS, respectively). Prediction errors were lowest in Alpine forests against GOSIF (MAE=130.89; RMSE=151.33 gC $m^{-2}$ $yr^{-1}$), while for MODIS the lowest MAE was found in the Apennine (215.98 gC $m^{-2}$ $yr^{-1}$) and the lowest RMSE in the Alpine ecoregion (271.96 gC $m^{-2}$ $yr^{-1}$).

Overall, 3D-CMCC-FEM successfully reproduced the magnitude, spatial variability, and temporal dynamics of GPP across Italian forests under present-day climatic conditions. Across all three climate forcings, model performance consistently followed the ranking ERA5-Land > UKESM1-0-LL > GFDL-ESM4, while agreement with GOSIF was systematically stronger than with MODIS. These results demonstrate that all three model configurations provide a robust basis for assessing future climate-driven changes in forest productivity.

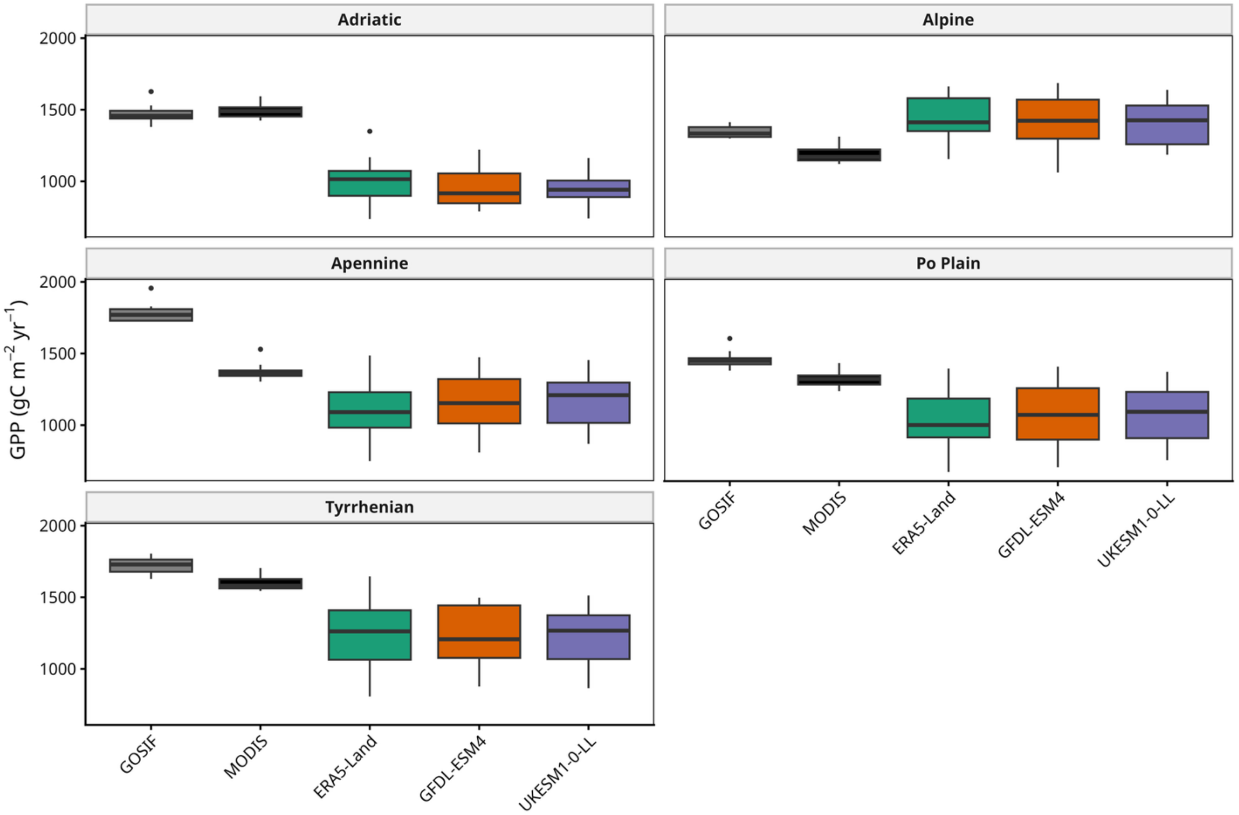


**Figure S1:** Distribution of GPP (gC $m^{-2}$ $yr^{-1}$) during the period 2005-2014 across five Italian ecoregions (i.e., Adriatic, Alpine, Apennine, Po Plain, and Tyrrhenian). Boxplots compare GPP estimates derived from GOSIF and MODIS products with 3D-CMCC-FEM simulations driven by ERA5-Land, GFDL-ESM4, and UKESM1-0-LL climate forcings. Boxes represent the interquartile range (IQR), the central line denotes the median, whiskers extend to 1.5 × IQR, and points indicate outliers.

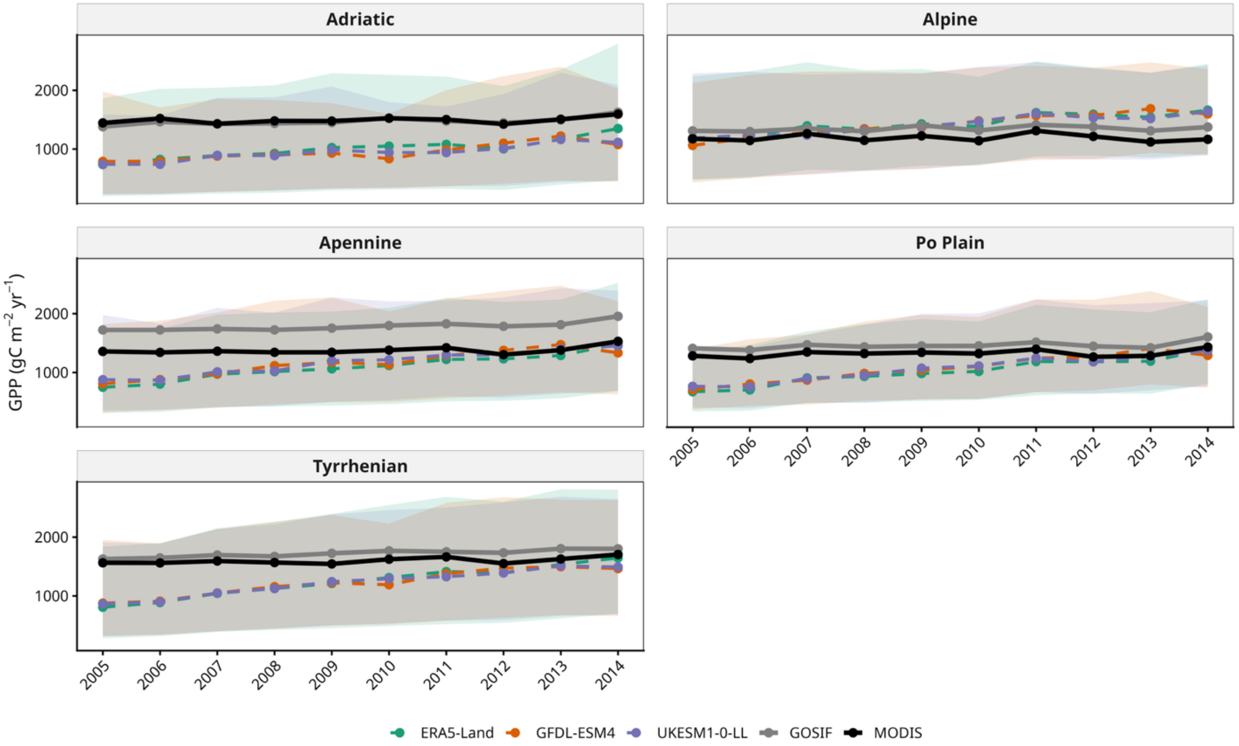


**Figure S2:** Temporal evolution of annual GPP (gC $m^{-2}$ $yr^{-1}$) from 2005 to 2014 across five Italian ecoregions (i.e., Adriatic, Alpine, Apennine, Po Plain, and Tyrrhenian). GPP simulated by 3D-CMCC-FEM driven by ERA5-Land, GFDL-ESM4, and UKESM1-0-LL climate forcings are compared with GPP estimates derived from GOSIF and MODIS RS-products. Solid lines represent annual regional mean GPP, while shaded areas indicate the 5th-95th percentile range across forest stands within each ecoregion.

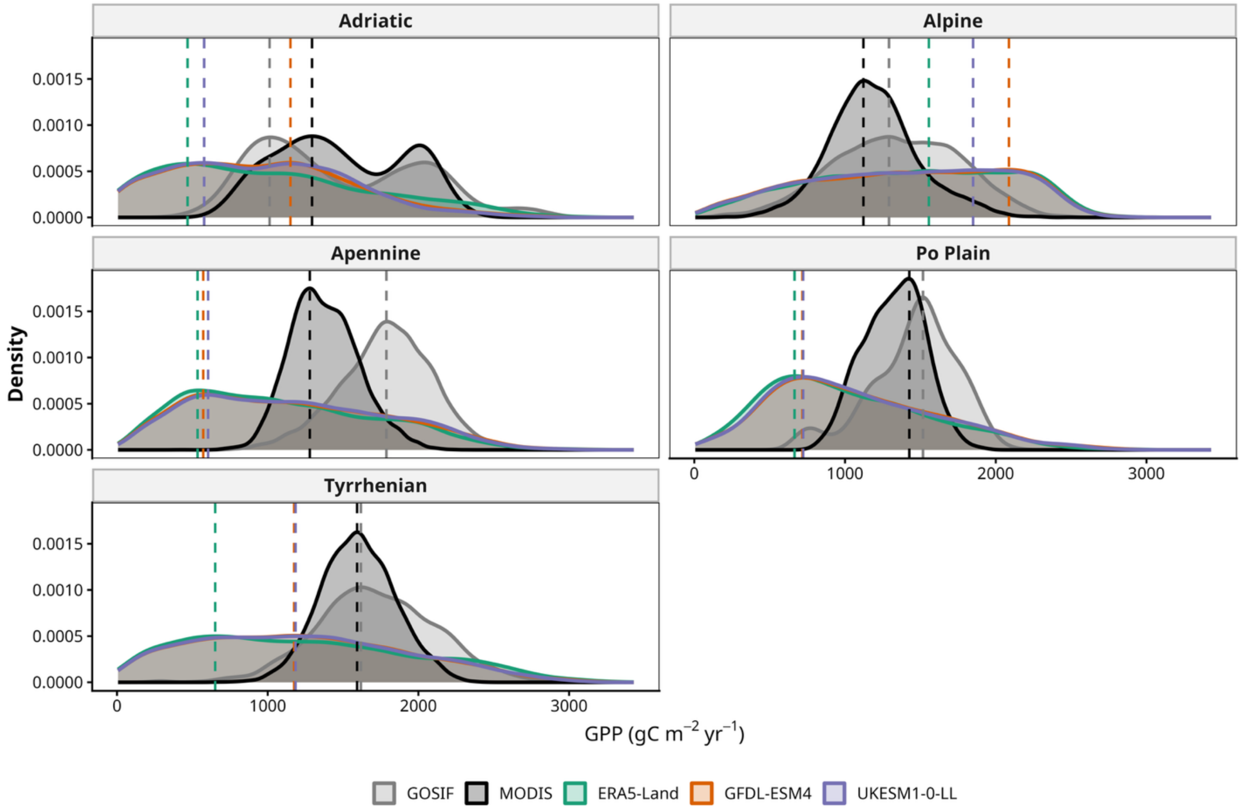


**Figure S3:** Probability Density Function (PDF) of GPP (gC $m^{-2}$ $yr^{-1}$) during the period 2005-2014 across five Italian ecoregions (i.e., Adriatic, Alpine, Apennine, Po Plain, and Tyrrhenian). Distributions are shown for 3D-CMCC-FEM simulations driven by ERA5-Land, GFDL-ESM4, and UKESM1-0-LL climate forcings, and compared with GOSIF- and MODIS-derived GPP estimates. Kernel density estimates were computed using annual GPP values from all forest stands and years within each ecoregion.

**Table S2:** Performance statistics of the 3D-CMCC-FEM model against GOSIF GPP for the historical period (2005-2014). Model performance is reported for simulations driven by ERA5-Land, GFDL-ESM4, and UKESM1-0-LL across the five Italian ecoregions (i.e., Adriatic, Alpine, Apennine, Po Plain, and Tyrrhenian). Statistical metrics include Bias (gC $m^{-2}$ $yr^{-1}$), mean absolute error (MAE; gC $m^{-2}$ $yr^{-1}$), root mean square error (RMSE; gC $m^{-2}$ $yr^{-1}$) and Pearson's correlation coefficient (R).

| FORCING CLIMATE | ECOREGION | Bias | MAE | RMSE | R |
|---|---|---|---|---|---|
| **ERA5-Land** | Adriatic | −467.96 | 467.96 | 481.68 | 0.88 |
| | Alpine | 86.71 | 135.76 | 162.7 | 0.68 |
| | Apennine | −690.22 | 690.22 | 707.61 | 0.89 |
| | Po Plain | −441.80 | 441.8 | 475.09 | 0.76 |
| | Tyrrhenian | −484.63 | 484.63 | 526.16 | 0.96 |
| **GFDL-ESM4** | Adriatic | −521.61 | 521.61 | 535.91 | 0.43 |
| | Alpine | 68.94 | 161.27 | 189.45 | 0.4 |
| | Apennine | −629.22 | 629.22 | 651.84 | 0.66 |
| | Po Plain | −387.93 | 387.93 | 435.19 | 0.5 |
| | Tyrrhenian | −500.35 | 500.35 | 527.97 | 0.88 |
| **UKESM1-0-LL** | Adriatic | −531.2 | 531.2 | 541.02 | 0.64 |
| | Alpine | 66.62 | 130.89 | 151.33 | 0.55 |
| | Apennine | −619.95 | 619.95 | 636.87 | 0.84 |
| | Po Plain | −396.59 | 396.59 | 430.44 | 0.7 |
| | Tyrrhenian | −503.14 | 503.14 | 528.86 | 0.96 |

**Table S3:** Performance statistics of the 3D-CMCC-FEM model against MODIS GPP for the historical period (2005-2014). Model performance is reported for simulations driven by ERA5-Land, GFDL-ESM4, and UKESM1-0-LL across the five Italian ecoregions (i.e., Adriatic, Alpine, Apennine, Po Plain, and Tyrrhenian). Statistical metrics include Bias (gC $m^{-2}$ $yr^{-1}$), mean absolute error (MAE; gC $m^{-2}$ $yr^{-1}$), root mean square error (RMSE; gC $m^{-2}$ $yr^{-1}$) and Pearson's correlation coefficient (R).

| FORCING CLIMATE | ECOREGION | Bias | MAE | RMSE | R |
|---|---|---|---|---|---|
| **ERA5-Land** | Adriatic | −484.9 | 484.9 | 504.15 | 0.68 |
| | Alpine | 241.88 | 245.83 | 286.5 | 0.33 |
| | Apennine | −281.77 | 281.77 | 335.53 | 0.63 |
| | Po Plain | −305.84 | 305.84 | 358.21 | 0.61 |
| | Tyrrhenian | −362.38 | 362.38 | 427.88 | 0.7 |
| **GFDL-ESM4** | Adriatic | −538.56 | 538.56 | 555.4 | 0.18 |
| | Alpine | 224.11 | 246.81 | 297.89 | 0.15 |
| | Apennine | −220.77 | 254.04 | 295.99 | 0.31 |
| | Po Plain | −251.98 | 276.65 | 325.65 | 0.36 |
| | Tyrrhenian | −378.11 | 378.11 | 425.83 | 0.52 |
| **UKESM1-0-LL** | Adriatic | −548.15 | 548.15 | 561.54 | 0.36 |
| | Alpine | 221.79 | 226.65 | 271.96 | 0.14 |
| | Apennine | −211.49 | 215.98 | 273.29 | 0.54 |
| | Po Plain | −260.63 | 260.63 | 314.85 | 0.58 |
| | Tyrrhenian | −380.9 | 380.9 | 426.59 | 0.59 |

##### 2.1.2. Evaluation against Italian NFI 2015

Although 3D-CMCC-FEM has previously been validated against the third Italian National Forest Inventory (INFI) (INFC, 2021) using high-resolution ERA5-derived meteorological forcing (Vangi et al., 2025b), the present analysis evaluates the model when driven by three alternative climate datasets (i.e., ERA5-Land, GFDL-ESM4, and UKESM1-0-LL). This additional assessment verifies that differences in climate forcing do not compromise the model's capability to reproduce the observed spatial variability of forest structural attributes before projecting future climate change impacts.

Simulated annual growing stock volume (GSV), aboveground biomass (AGB), and carbon stock (CS) for the inventory year (2015) were aggregated across the five Italian ecoregions and compared with the corresponding estimates from the INFI. The PDF shown in Fig. S4 closely matched those derived from the INFI, reproducing both the central tendency and the overall shape of the observed distributions. For GSV, the highest probability densities occurred between approximately 100 and 300 $m^3$ $ha^{-1}$, with long right tails extending beyond 500-800 $m^3$ $ha^{-1}$, particularly in the Alpine and Tyrrhenian ecoregions. AGB exhibited a similar pattern, with most stands ranging between 50 and 150 Mg $ha^{-1}$, while only a limited fraction exceeded 300 Mg $ha^{-1}$. Likewise, CS was primarily distributed between 30 and 100 MgC $ha^{-1}$, with relatively few stands exceeding 200 MgC $ha^{-1}$. Among the three climate forcings used to drive 3D-CMCC-FEM, ERA5-Land produced distributions that most closely resembled the observations, whereas GFDL-ESM4 and UKESM1-0-LL yielded nearly overlapping curves. These results were further confirmed by the comparison of ecoregion-averaged means (± standard deviation) (Fig. S5), which showed close agreement between simulations and INFI observations. ERA5-Land again provided the closest agreement with the observed means, while GFDL-ESM4 and UKESM1-0-LL produced nearly identical estimates.

The quantitative performance metrics provided additional evidence of the capability of 3D-CMCC-FEM to reproduce the spatial patterns of forest structural attributes across Italy. Taylor diagrams further confirmed the model's ability to reproduce the observed spatial variability of forest structural attributes. R ranged from 0.52 to 0.86 for GSV, 0.54 to 0.83 for AGB, and 0.52 to 0.81 for CS, with ERA5-Land consistently achieving the highest agreement with the INFI values (Fig. S6). Biases indicated a systematic underestimation of GSV (−59.37 to −6.97 $m^3$ $ha^{-1}$) and AGB (−31.81 to 4.94 Mg $ha^{-1}$), whereas CS was generally overestimated (0.33 to 23.72 MgC $ha^{-1}$). MAE varied between 58.90 and 97.74 $m^3$ $ha^{-1}$ for GSV, 41.48 and 55.31 Mg $ha^{-1}$ for AGB, and 24.62 and 36.13 MgC $ha^{-1}$ for CS, while RMSE ranged from 87.32 to 150.82 $m^3$ $ha^{-1}$, 65.49 to 99.54 Mg $ha^{-1}$, and 33.42 to 48.36 MgC $ha^{-1}$, respectively (Tables S4-S6).

In general, model performance was highest for GSV, followed by AGB and CS. ERA5-Land consistently provided the best agreement with the INFI across all structural attributes, whereas differences between GFDL-ESM4 and UKESM1-0-LL remained negligible, indicating that the choice of climate forcing had only a limited influence on the representation of present-day forest structural attributes.

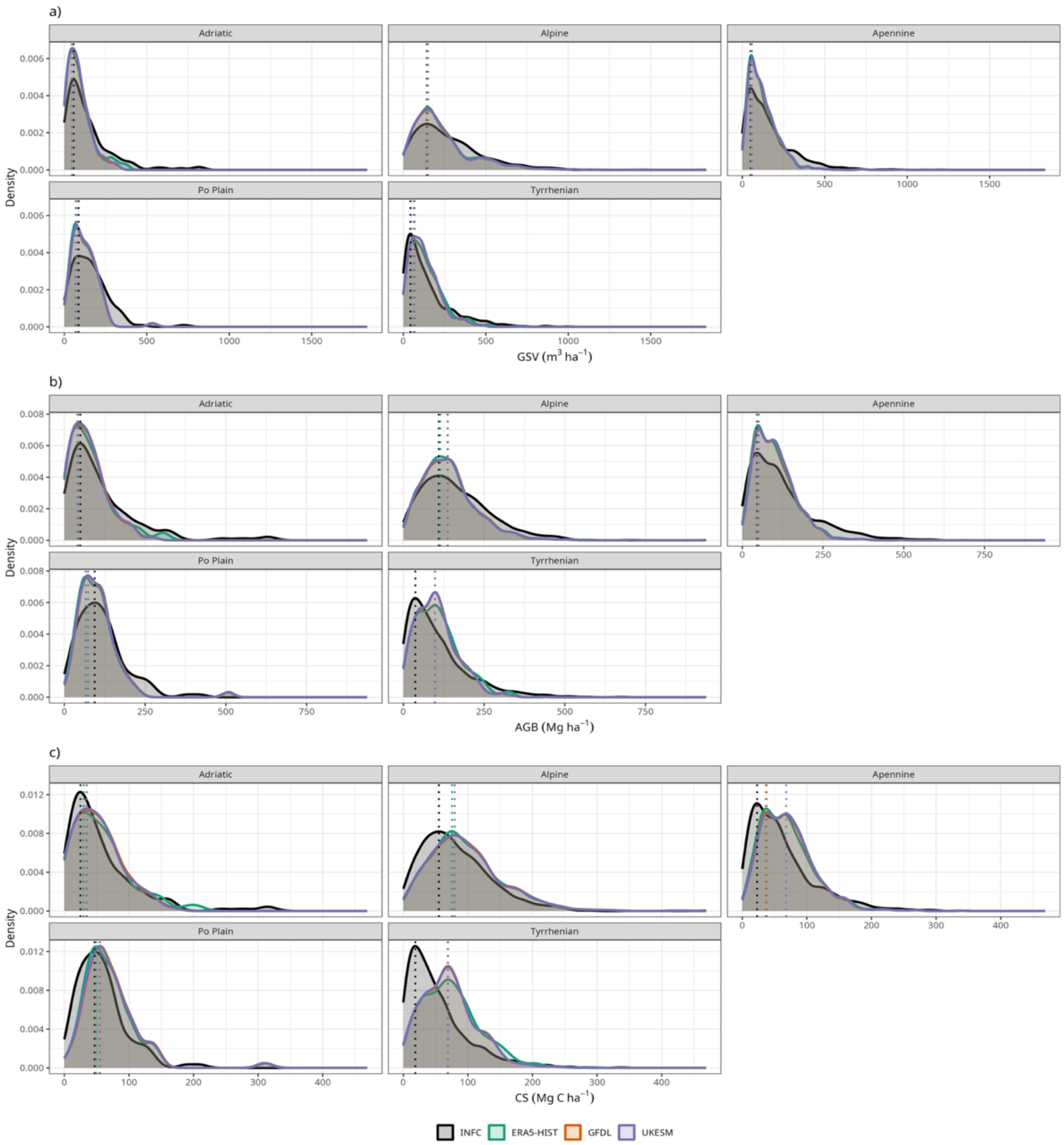


**Figure S4:** Comparison of observed and simulated distributions of (a) growing stock volume (GSV; $m^{-3}$ $ha^{-1}$), (b) above ground biomass (AGB; Mg $ha^{-1}$), and (c) carbon stock (CS; MgC $ha^{-1}$) for the five Italian forest ecoregions (i.e., Adriatic, Alpine, Apennine, Po Plain, and Tyrrhenian). Black lines represent observations from the INFC, whereas colored lines show simulations from the 3D-CMCC-FEM model driven by ERA5-Land, GFDL-ESM4, and UKESM1-0-LL climate datasets. Vertical dashed lines indicate the corresponding mean values of each distribution. Results refer to annual stand-level values for 2015.

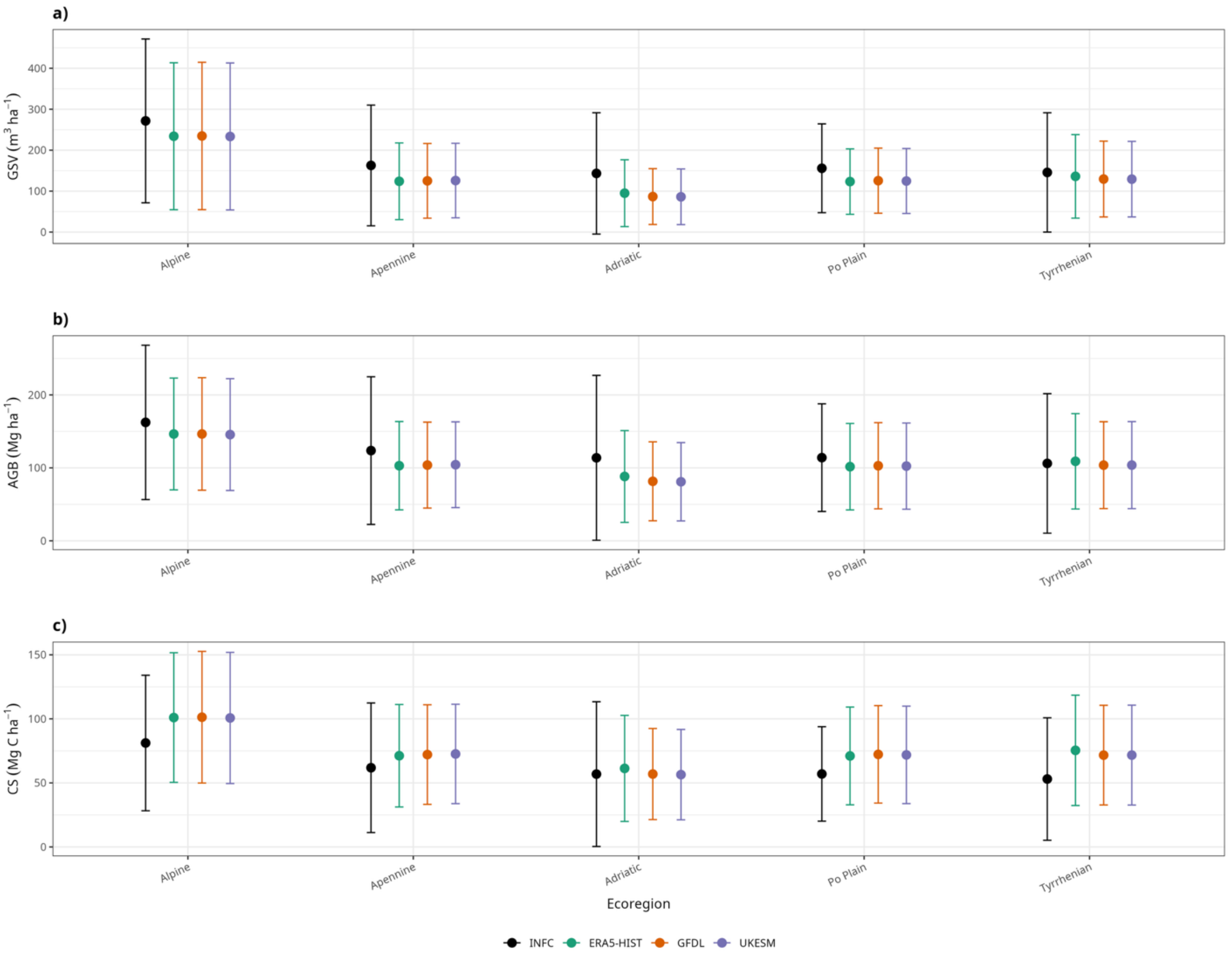


**Figure S5:** Mean ± standard deviation of (a) Growing Stock Volume (GSV; $m^{-3}$ $ha^{-1}$), (b) Above Ground Biomass (AGB; Mg $ha^{-1}$), and (c) Carbon Stock (CS; MgC $ha^{-1}$) for the five Italian forest ecoregions (i.e., Alpine, Apennine, Adriatic, Po Plain, and Tyrrhenian). Black symbols represent observations from the third Italian National Forest Inventory (INFC, 2021), while colored symbols represent simulations obtained with the 3D-CMCC-FEM model forced by ERA5-Land, GFDL-ESM4, and UKESM1-0-LL climate datasets. Values correspond to annual means for 2015 aggregated at the ecoregion level.

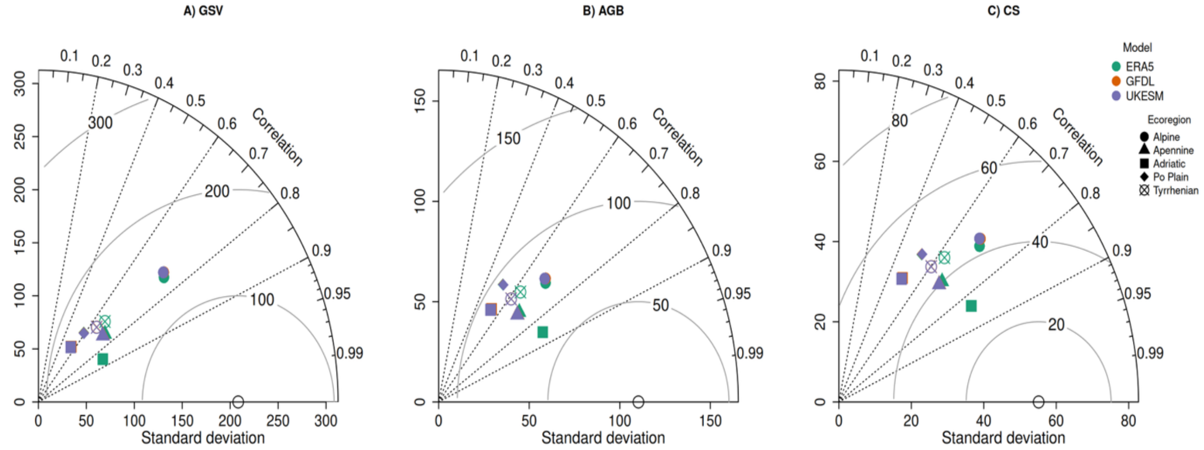


**Figure S6:** Taylor diagrams comparing simulated and observed values of (A) Growing Stock Volume (GSV), (B) Aboveground Biomass (AGB), and (C) Carbon Stock (CS) across the five Italian forest ecoregions. Colored symbols represent the three climate datasets used to drive the simulations (i.e., ERA5, GFDL, and UKESM), while symbol shapes identify the ecoregions (i.e., Alpine, Apennine, Adriatic, Po Plain, and Tyrrhenian). Results are based on the average annual values for 2015. The radial distance from the origin indicates the model standard deviation (SD), the angular coordinate represents the Pearson's correlation coefficient (R) with observations, and the semicircles denote the centered root mean square difference (RMSD). The open circle on the x-axis indicates the observed standard deviation (SD).

**Table S4:** Performance statistics of the 3D-CMCC-FEM model against the Growing Stock Volume (GSV) estimates derived from the third Italian National Forest Inventory (INFC, 2021). Model performance is reported for simulations driven by ERA5-Land, GFDL-ESM4, and UKESM1-0-LL across the five Italian ecoregions (i.e., Adriatic, Alpine, Apennine, Po Plain, and Tyrrhenian). Statistical metrics include Bias ($m^{-3}$ $ha^{-1}$ $yr^{-1}$), mean absolute error (MAE; $m^{-3}$ $ha^{-1}$ $yr^{-1}$), root mean square error (RMSE; $m^{-3}$ $ha^{-1}$ $yr^{-1}$) and Pearson's correlation coefficient (R).

| FORCING CLIMATE | ECOREGION | Bias | MAE | RMSE | R |
|---|---|---|---|---|---|
| **ERA5-Land** | Adriatic | −49.57 | 68.68 | 114.57 | 0.86 |
| | Alpine | −27.72 | 96.09 | 143.49 | 0.74 |
| | Apennine | −39.31 | 72.46 | 110.9 | 0.73 |
| | Po Plain | −27.06 | 59.83 | 88.22 | 0.59 |
| | Tyrrhenian | −6.97 | 70.98 | 110.27 | 0.68 |
| **GFDL-ESM4** | Adriatic | −58.73 | 78.04 | 150.24 | 0.55 |
| | Alpine | −25.85 | 97.06 | 147 | 0.73 |
| | Apennine | −38.19 | 72.38 | 111.1 | 0.73 |
| | Po Plain | −24.89 | 59.06 | 87.32 | 0.59 |
| | Tyrrhenian | −13.6 | 72.55 | 113.76 | 0.65 |
| **UKESM1-0-LL** | Adriatic | −59.37 | 78.28 | 150.82 | 0.55 |
| | Alpine | −27.08 | 97.74 | 147.44 | 0.73 |
| | Apennine | −37.49 | 72.18 | 110.81 | 0.73 |
| | Po Plain | −25.77 | 58.9 | 87.34 | 0.59 |
| | Tyrrhenian | −13.82 | 72.61 | 113.87 | 0.65 |

**Table S5:** Performance statistics of the 3D-CMCC-FEM model against Aboveground Biomass (AGB) estimates derived from the third Italian National Forest Inventory (INFC, 2021). Model performance is reported for simulations driven by ERA5-Land, GFDL-ESM4, and UKESM1-0-LL across the five Italian ecoregions (i.e., Adriatic, Alpine, Apennine, Po Plain, and Tyrrhenian). Statistical metrics include Bias (Mg $ha^{-1}$ $yr^{-1}$), mean absolute error (MAE; Mg $ha^{-1}$ $yr^{-1}$), root mean square error (RMSE; Mg $ha^{-1}$ $yr^{-1}$) and Pearson's correlation coefficient (R).

| FORCING CLIMATE | ECOREGION | Bias | MAE | RMSE | R |
|---|---|---|---|---|---|
| **ERA5-Land** | Adriatic | −23.89 | 46.51 | 73.37 | 0.83 |
| | Alpine | −15.88 | 54.29 | 76.38 | 0.71 |
| | Apennine | −20.04 | 51.56 | 76.63 | 0.69 |
| | Po Plain | −12.3 | 41.97 | 66.18 | 0.54 |
| | Tyrrhenian | 4.94 | 53.24 | 76.4 | 0.61 |
| **GFDL-ESM4** | Adriatic | −31.18 | 52.4 | 99.15 | 0.56 |
| | Alpine | −15.57 | 54.87 | 77.75 | 0.69 |
| | Apennine | −19.18 | 51.67 | 76.64 | 0.69 |
| | Po Plain | −10.33 | 41.49 | 65.49 | 0.54 |
| | Tyrrhenian | −0.46 | 53.75 | 78.54 | 0.57 |
| **UKESM1-0-LL** | Adriatic | −31.81 | 52.64 | 99.54 | 0.55 |
| | Alpine | −16.45 | 55.31 | 78.32 | 0.69 |
| | Apennine | −18.62 | 51.59 | 76.49 | 0.69 |
| | Po Plain | −10.81 | 41.48 | 65.56 | 0.54 |
| | Tyrrhenian | −0.38 | 53.56 | 78.15 | 0.58 |

**Table S6:** Performance statistics of the 3D-CMCC-FEM model against the Carbon Stock (CS) estimates derived from the third Italian National Forest Inventory (INFC, 2021). Model performance is reported for simulations driven by ERA5-Land, GFDL-ESM4, and UKESM1-0-LL across the five Italian ecoregions (i.e., Adriatic, Alpine, Apennine, Po Plain, and Tyrrhenian). Statistical metrics include Bias (MgC $ha^{-1}$ $yr^{-1}$), mean absolute error (MAE; MgC $ha^{-1}$ $yr^{-1}$), root mean square error (RMSE; MgC $ha^{-1}$ $yr^{-1}$) and Pearson's correlation coefficient (R).

| FORCING CLIMATE | ECOREGION | Bias | MAE | RMSE | R |
|---|---|---|---|---|---|
| **ERA5-Land** | Adriatic | 5.6 | 24.62 | 33.42 | 0.81 |
| | Alpine | 19.9 | 33.47 | 44.09 | 0.71 |
| | Apennine | 9.79 | 28.85 | 39.3 | 0.67 |
| | Po Plain | 14.24 | 25.27 | 38.21 | 0.56 |
| | Tyrrhenian | 23.72 | 36.13 | 47.05 | 0.61 |
| **GFDL-ESM4** | Adriatic | 0.82 | 28.22 | 48.31 | 0.52 |
| | Alpine | 20.34 | 34.32 | 45.48 | 0.69 |
| | Apennine | 10.71 | 29.35 | 39.52 | 0.67 |
| | Po Plain | 15.79 | 25.89 | 38.59 | 0.56 |
| | Tyrrhenian | 19.91 | 34.98 | 45.77 | 0.57 |
| **UKESM1-0-LL** | Adriatic | 0.33 | 28.16 | 48.36 | 0.52 |
| | Alpine | 19.77 | 34.3 | 45.45 | 0.69 |
| | Apennine | 11.23 | 29.54 | 39.63 | 0.67 |
| | Po Plain | 15.39 | 25.67 | 38.43 | 0.56 |
| | Tyrrhenian | 19.96 | 34.92 | 45.63 | 0.57 |

#### 2.2. Climate change projections

##### 2.2.1. Temporal patterns of carbon dynamics across Italian ecoregions (2015-2100)

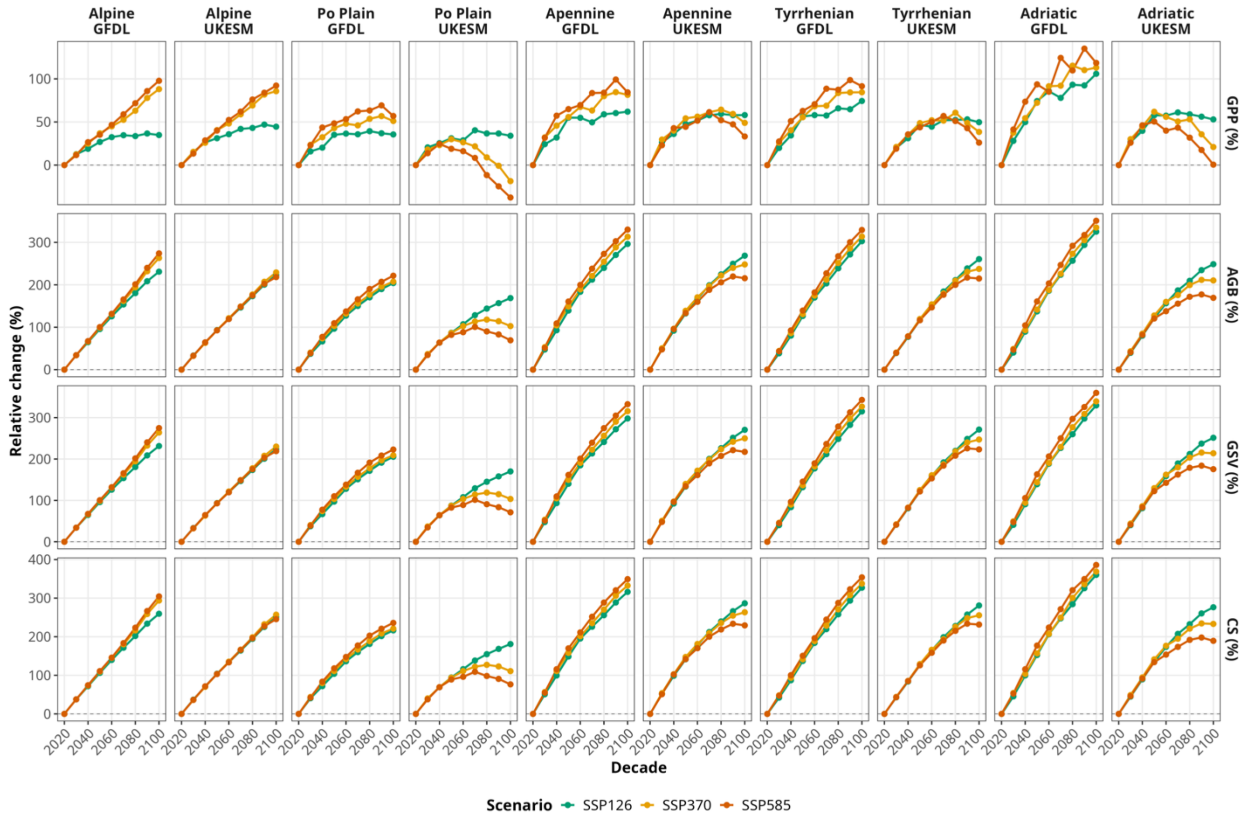


**Figure S7:** Relative change in forest productivity and carbon stocks compared with the 2020 decade across Italian ecoregions (i.e., Alpine, Apennine, Adriatic, Po Plain, and Tyrrhenian). Relative (%) changes in of GPP, Aboveground Biomass (AGB), Growing Stock Volume (GSV) and Carbon Stock (CS) with respect to the 2020 decade, calculated for successive decades until 2100 (i.e. 2015-2025, 2026-2035 etc.). Results are shown for each Italian ecoregion, climate scenario, and ESM (GFDL-ESM4 and UKESM1-0-LL).

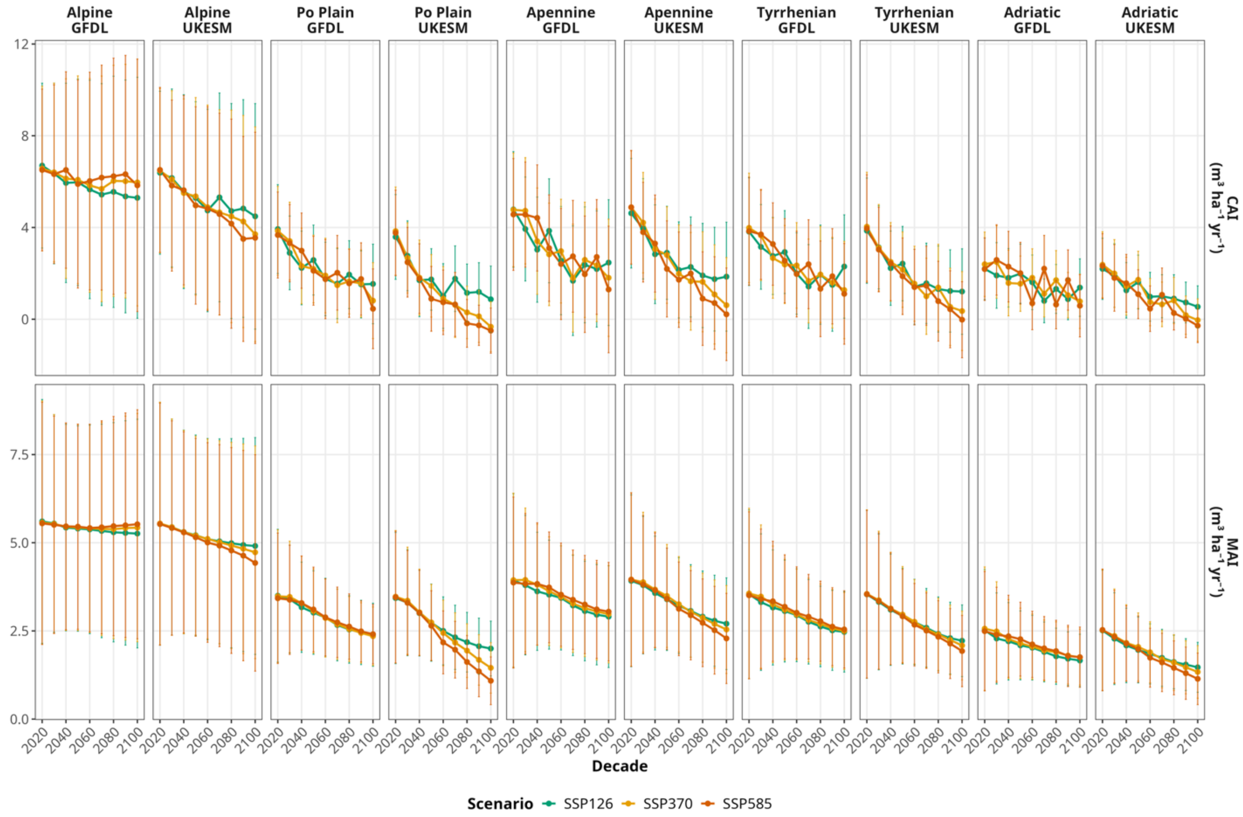


**Figure S8:** Decadal changes in stand growth indices. Mean decadal values (±1 standard deviation) of Current Annual Increment (CAI; $m^3\ ha^{-1}\ yr^{-1}$) and Mean Annual Increment (MAI; $m^3\ ha^{-1}\ yr^{-1}$) for the five Italian ecoregions (i.e., Alpine, Apennine, Adriatic, Po Plain, and Tyrrhenian) under SSP126, SSP370, and SSP585 scenario, simulated with the GFDL-ESM4, and UKESM1-0-LL climate forcings.

##### 2.2.2. Temporal patterns of carbon dynamics across forest types (2015-2100)

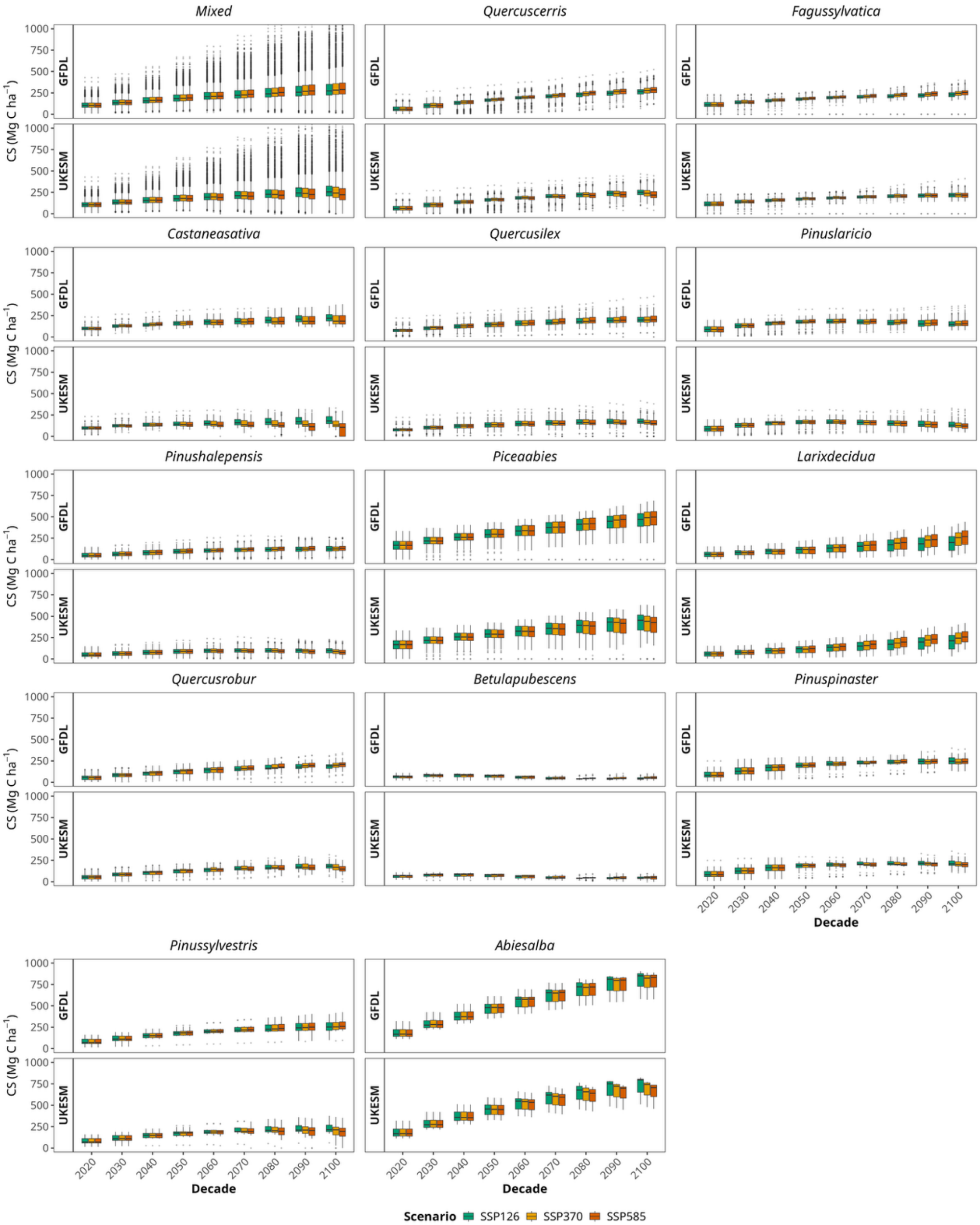


**Figure S9:** Temporal evolution of Carbon Stock (CS, Mg C ha$^{-1}$) across the dominant forest types in Italy under the SSP126, SSP370, and SSP585 climate scenarios. Results are presented separately for the GFDL and UKESM climate forcings. Boxplots represent the spatial variability among simulated forest stands within each forest type, while dashed lines connect the decadal means. Results are shown for successive decades from 2020 to 2100. Boxplots summarize the distribution across all INFI stands belonging to each forest type; the central line denotes the median, boxes the interquartile range, whiskers extend to 1.5 × IQR, and points represent outliers.

##### 2.2.3. Temporal variability of forest productivity across Italian ecoregions (2015-2100)

**Table S7:** Ecoregion Mean Coefficient of Variation (CV, %) of GPP for the 2020 and 2100 decades and the corresponding relative change (ΔCV, %) across the five Italian ecoregions (i.e., Alpine, Apennine, Adriatic, Po Plain, and Tyrrhenian) under the SSP126, SSP370, and SSP585 climate scenarios. Results are reported separately for simulations driven by the GFDL-ESM4 and UKESM1-0-LL climate forcings. Positive ΔCV values indicate an increase in productivity variability relative to the 2020 baseline, whereas negative values indicate a decrease.

| Ecoregion | UKESM1-0-LL | | | GFDL-ESM4 | | |
|---|---|---|---|---|---|---|
| | $CV_{2020}$ | $CV_{2100}$ | Delta CV | $CV_{2020}$ | $CV_{2100}$ | Delta CV |
| **SSP126** | | | | | | |
| **Alpine** | 8.43 | 5.47 | −35.1 | 7.05 | 3.8 | −46.1 |
| **Apennine** | 12.37 | 7.13 | −42.3 | 11.34 | 4.31 | −62 |
| **Adriatic** | 13.34 | 14.2 | 6.4 | 13.77 | 9.14 | −33.7 |
| **Po Plain** | 9.96 | 6.57 | −34 | 8.54 | 3.58 | −58.1 |
| **Tyrrhenian** | 9.85 | 9.38 | −4.7 | 9.8 | 3.81 | −61.1 |
| **SSP370** | | | | | | |
| **Alpine** | 9.36 | 7.09 | −24.3 | 7.74 | 3.44 | −55.5 |
| **Apennine** | 13.25 | 12.05 | −9.1 | 10.72 | 9.36 | −12.7 |
| **Adriatic** | 13.08 | 11.6 | −11.3 | 13.06 | 14.99 | 14.8 |
| **Po Plain** | 11.09 | 19.3 | 74 | 9.07 | 7.48 | −17.6 |
| **Tyrrhenian** | 10.4 | 11.85 | 14 | 9.53 | 8.72 | −8.6 |
| **SSP585** | | | | | | |
| **Alpine** | 9.22 | 8.64 | −6.3 | 8.8 | 3.92 | −55.5 |
| **Apennine** | 11.69 | 9.3 | −20.4 | 12.54 | 8.98 | −28.4 |
| **Adriatic** | 12.82 | 8.62 | −32.8 | 14.49 | 13.26 | −8.5 |
| **Po Plain** | 10.17 | 23.62 | 132.2 | 11.61 | 7.64 | −34.2 |
| **Tyrrhenian** | 9.86 | 10.63 | 7.8 | 10.78 | 9.1 | −15.5 |

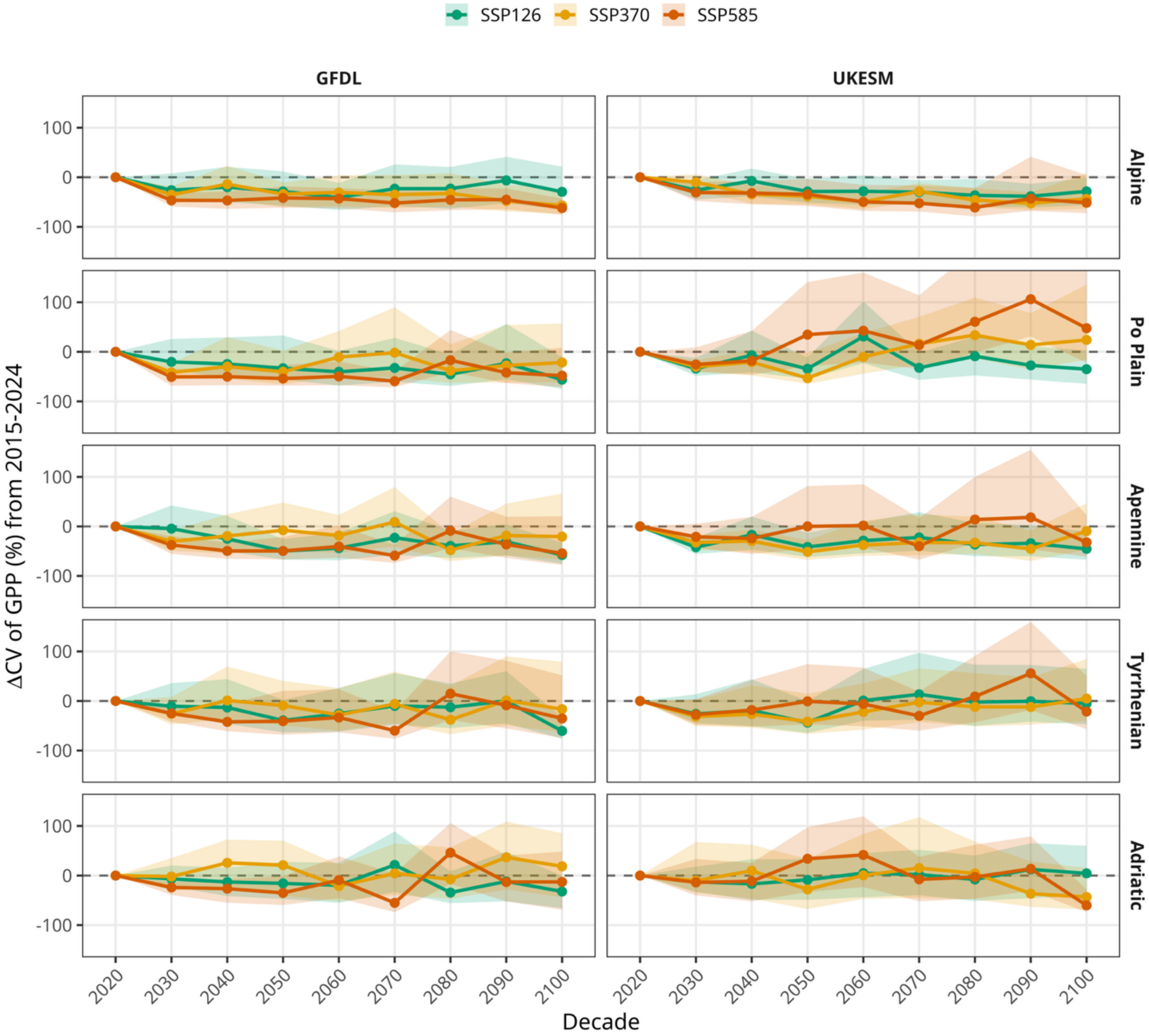


**Figure S10:** Relative change in the Coefficient of Variation (CV, %) of GPP with respect to the 2020 decade, calculated for successive decades until 2100 (i.e. 2015-2025, 2026-2035 etc.) across the five Italian ecoregions (i.e., Alpine, Apennine, Adriatic, Po Plain, and Tyrrhenian) under SSP126, SSP370, and SSP585 scenario. Shaded areas represent ±1 standard deviation among forest stands. Results are shown separately for the GFDL-ESM4 and UKESM1-0-LL climate forcings.

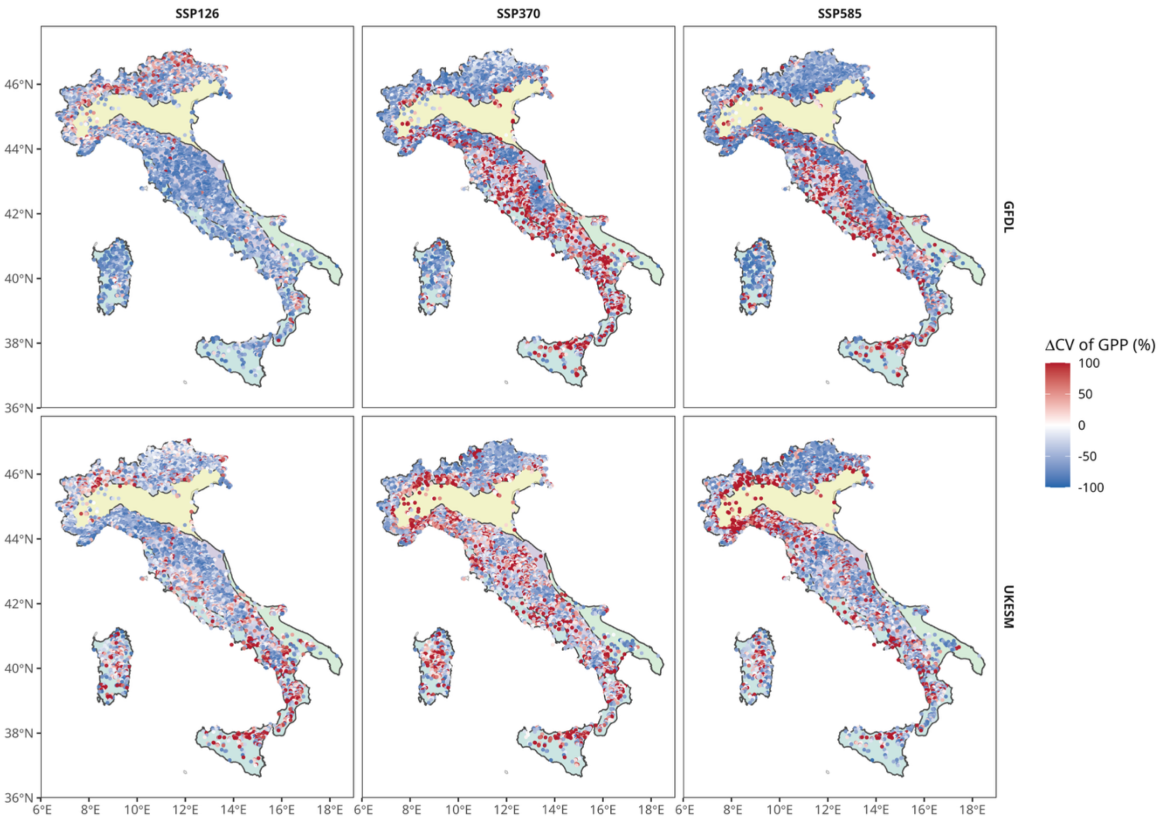


**Figure S11:** Spatial distribution of changes in GPP variability. Relative change in the Coefficient of Variation (CV, %) of GPP between the 2100 and 2020 decades for individual forest stands across Italy. Maps are shown for the SSP126, SSP370, and SSP585 scenarios (columns) and for the GFDL-ESM4 and UKESM1-0-LL ESMs (rows). Red colors indicate an increase in interannual variability, whereas blue colors indicate a decrease relative to the baseline period. The colored areas represent the ecoregions following the Fig. 1.

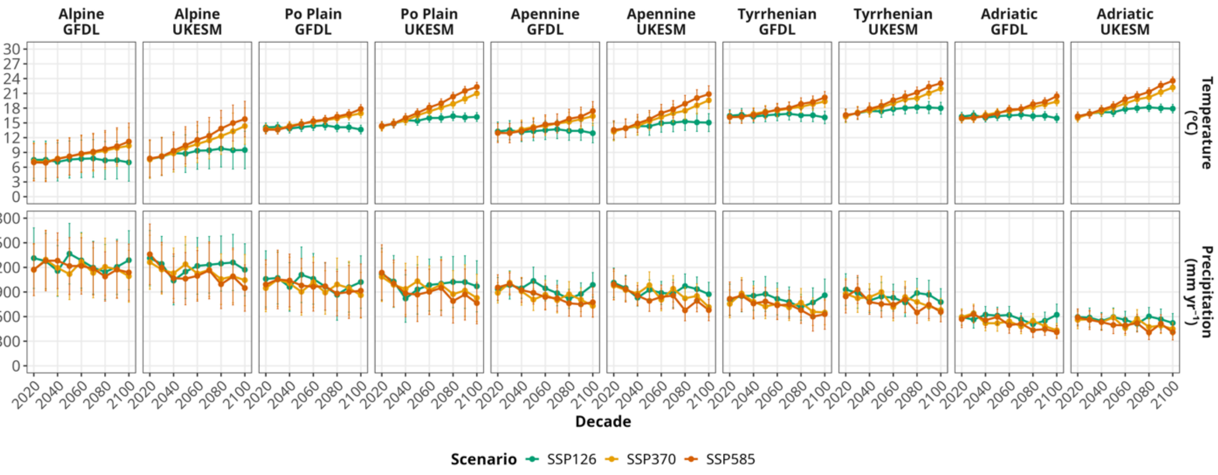


**Figure S12:** Projected changes in mean decadal temperature (°C) and precipitation (mm yr$^{-1}$) (±1 standard deviation) from 2020 to 2100 across the five Italian ecoregions (i.e., Alpine, Po Plain, Apennine, Tyrrhenian, and Adriatic) under the GFDL-ESM4 and UKESM1-0-LL ESMs and three SSP scenarios (i.e., SSP1-2.6, SSP3-7.0, and SSP5-8.5).
